# Geometric influences on the regional organization of the mammalian brain

**DOI:** 10.1101/2025.01.30.635820

**Authors:** James C. Pang, Peter A. Robinson, Kevin M. Aquino, Priscila T. Levi, Alexander Holmes, Marija Markicevic, Xilin Shen, Francesca Mandino, Thomas Funck, Nicola Palomero-Gallagher, Ru Kong, B.T. Thomas Yeo, Jeggan Tiego, Mark A. Bellgrove, R Todd Constable, Evelyn Lake, Michael Breakspear, Alex Fornito

## Abstract

The mammalian brain comprises anatomically and functionally distinct regions, yet the principles governing their organization across scales remain unclear. Efforts to map regional boundaries use expert judgement or data-driven clustering of functional, connectional, and/or architectonic properties, but these approaches are often descriptive, have limited generalizability, and do not elucidate generative mechanisms. Here, we develop a geometrically constrained framework for the hierarchical, multiscale regional organization of neocortical and non-neocortical structures. The framework yields parcels at any resolution with high internal homogeneity across hundreds of anatomical, functional, cellular, and molecular properties in humans, macaques, marmosets, and mice, and it generalizes to understudied mammalian species. Finally, we demonstrate that the framework captures the essence of a reaction-diffusion mechanism in which brain geometry shapes the spatial expression of putative patterning molecules that establish a blueprint of regional identity during development. Our findings point to a conserved, universal influence of geometry on mammalian brain organization.

## Introduction

Over 100 years ago, Brodmann famously used variations in cytoarchitecture to subdivide, or parcellate, the human cerebral cortex into 43 discrete areas^1^, which he considered to be “specific morphological organs” (p. 251), each representing an “exclusive individual function that is different from those of other organs” (pp. 251–252). His proposal that the brain is composed of anatomically and functionally specialized areas has fundamentally influenced a century of subsequent research^2,3^, such that the prevailing orthodoxy in contemporary neuroscience is that coordinated behaviour arises from interactions between these discrete, specialized areas^4,5^.

Despite the popularity of this paradigm, accurate and reliable delineation of areal boundaries has been challenging. Different anatomists have proposed cytoarchitectonic parcellations that differ in both the number of areas defined and the anatomical locations of their boundaries^6^, leading some to question their validity and utility, particularly beyond sharply delimited primary sensorimotor cortices^7–9^. More recent work has leveraged advances in non-invasive techniques, such as magnetic resonance imaging (MRI), to statistically cluster brain locations according to similarities in microstructural, anatomical, connectivity, and/or functional profiles^10–13^. This work culminated in a flagship study of the Human Connectome Project (HCP)^14^, which synthesized multimodal measures to define 180 discrete regions per cortical hemisphere^2^––a >4-fold increase in the number of regions over Brodmann’s original map.

This body of research has been foundational to our understanding of regional specialization in the brain, but it also highlights that such organization can be conceptualized in different ways^3^. Accordingly, for present purposes we draw a distinction between *areas* and *parcels.* Areas refer to local, contiguous patches of neural tissue with distinct cellular, molecular, and/or functional specialization classically identified through histology (e.g., V1, MT). Parcels refer to contiguous patches of tissue defined through data-driven clustering of high-dimensional data, as commonly (but not exclusively) measured with neuroimaging; for example, by characterizing how distinct brain locations are functionally coupled through time^10^. We use the terms *regions* and *regional organization* to refer more broadly to the way in which the local specialization of any anatomical or functional property is organized, regardless of whether it is defined in terms of areas or parcels. Areas and parcels can overlap, but they can also diverge because they are defined in different ways and prioritize different properties for defining regional boundaries. For example, the HCP parcellation was derived by calculating spatial gradients in multimodal brain imaging data as a starting point for defining brain parcels, before relying on manual intervention in specific cases to select parcel boundaries that fit prior anatomical knowledge of areal divisions^2^.

Discrepancies between areal and parcel boundaries reflect the broader challenge that no single definitive map can fully capture the brain’s complex regional organization; instead, the optimal choice depends on the question asked of the data. Indeed, classical approaches to dividing the brain into areas or parcels make several limiting assumptions. First, they assume sharp transitions in brain structure and/or function, even though the primate cortex often shows graded architectonic transitions^1,7,15–17^. Second, borders defined according to different properties (e.g., cytoarchitecture, myeloarchitecture, brain activity) do not always coincide^6,18^, raising questions about which specific property should be prioritized; even different measures of similar properties can diverge, given recent evidence in marmoset cortex that sharp spatial transitions of transcriptomically defined cell types align with classical cytoarchitectonic boundaries only 30–40% of the time^19^. Third, functional parcel boundaries can shift with changing cognitive demands^20–22^. Fourth, cytoarchitectonically defined areas contain neurons with diverse functional responses, meaning that activity maps often show poor correspondence with cytoarchitecture and that the neural correlates of behaviour often transcend areal boundaries^23–27^. Fifth, most areal and parcel divisions are typically defined at a single spatial resolution when, in fact, the brain shows an intrinsically multiscale organization^28–30^ that spans supra-regional (e.g., broad cortical “types” of laminar differentiation^31–35^ and distributed, large-scale networks^36–38^) and infra-regional (e.g., columns and hypercolumns^39^) scales. Accordingly, multiple, spatially overlapping modes of structural and functional organization are evident even within regions with well-defined borders, such as V1^40^, or those with clear subnuclear structure, such as the thalamus^41^. Finally, most approaches to brain regionalization are primarily descriptive, identifying areas or parcels by expert judgment or statistical clustering of patterns in data without direct links to an underlying generative mechanism (for exceptions, see refs^42–46^). Together, these considerations suggest that traditional maps of areas or parcels describe important, yet limited features of regional organization that do not shed light on underlying generative mechanisms or generalize across different contexts, resolutions, individuals, brain structures, and species.

Here, we address these limitations by adopting a different approach inspired by growing evidence that the geometry of the brain fundamentally constrains its structural and functional organization^47–51^, including the spatial expression profiles of molecules that play an essential role in regional patterning^52,53^ (see also next section). We thus develop a simple framework that captures the essence of key geometric and physical constraints on the regional patterning of the mammalian brain, allowing us to characterize the hierarchical, multiscale organization of neocortical and non-neocortical structures. Rather than clustering high-dimensional brain features into a statistically optimal map of putative areas or parcels, our approach only requires a model of the geometry of a brain structure as input, allowing us to assess the degree to which regional organization can be explained by the geometry of a brain structure alone. The framework uses geometric eigenmodes^47^ to recursively divide brain structures into nested parcels, providing an operational readout to test the hypothesis that geometry supplies an initial scaffold for multiscale organization. Critically, the method provides a direct mapping between discrete and more graded accounts of brain architecture, whilst also offering an intrinsically multiscale description of regional organization. We show that these geometry-derived parcels are internally homogeneous across hundreds of anatomical, functional, cellular, and molecular properties measured in humans, macaques, marmosets, and mice, often matching or exceeding the homogeneity of parcels defined by existing benchmark atlases. We further show that our approach aligns naturally with a classical reaction-diffusion mechanism of patterning substances that play an important, and perhaps a minimally sufficient role, in shaping the initial blueprint of regional organization in the brain.

### A geometrically constrained framework for the brain’s hierarchical regional organization

Early regionalization of brain structures is shaped by morphogens, which are secreted from specific patterning centers and diffuse through the developing cortical primordium along spatially continuous gradients^54–56^. For instance, in the midbrain and hindbrain, highly conserved gradients of *Hox* genes are sufficient to sharply delineate distinct expression domains, called prosomeres, that give rise to subdivisions of the diencephalon and secondary prosencephalon^57^. In the cortex, morphogens, such as fibroblast growth factor 8 (FGF8), Wnt, and Sonic Hedgehog (Shh), trigger the subsequent expression of distinct transcription factors (e.g., *Pax6*, *Emx2*, *Couptf1*) that further drive areal specification^58–64^. The superposition of these expression gradients spatially organizes neurogenic and other developmental dynamics, creating a rudimentary blueprint of areal identity that is gradually refined by thalamic innervation and activity-dependent mechanisms^52,63,65–69^.

The expression gradients formed by morphogens and other patterning factors predominantly align with the rostrocaudal, dorsoventral, and mediolateral axes of the developing telencephalon^52,65,70–73^. As such, they correspond to the cardinal axes of geometric variation in brain structure, which are formally described by the low-order geometric eigenmodes of the cortex derived from a mathematical decomposition of cortical geometry that is comparable to Fourier decomposition^47,74–76^ (see details below). The equivalence between the expression gradients and geometric eigenmodes arises because the geometry of a medium shapes the spatial patterns that arise from the interactions and diffusion of any molecules within it, as dictated by Turing’s classical reaction-diffusion equations^56,77^. The low-order eigenmodes are the least physically stable in generic reaction-diffusion systems and are therefore the first patterns to be expressed^78^. The preponderance of such patterns in early expression gradients of the developing brain^52^ thus arises from the physical constraints that geometry imposes on molecular dynamics.

We leverage this constraining effect of geometry to define a hierarchical framework for approximating the brain’s regional organization, capturing the essential features of a hypothesized set of reaction-diffusion processes, without having to specify the full set of biophysical details of the molecular interactions involved. We begin by describing the application of our framework to the left hemisphere of the human neocortex, the surface geometry of which is approximated by a triangular mesh (with 32,492 vertices) derived from a population-average template extracted from T1-weighted (T1w) MRI (Fig. 1A, top left). First, we calculate the surface’s geometric eigenmodes by solving the Helmholtz equation,

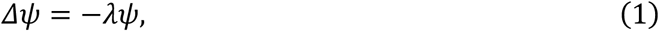

where *Δ* is the Laplace-Beltrami operator (LBO), which accounts for the spatial relationships between vertices on the surface mesh (see ‘Derivation of geometric eigenmodes’ in Methods), and *ψ* = {*ψ*_1_, *ψ*_2_, …} is the family of geometric eigenmodes with corresponding eigenvalues, *λ* = {*λ*_1_, *λ*_2_, …}. The eigenvalues are arranged in increasing order and correspond to the effective spatial wavelength of each eigenmode, with *λ*_1_ ≈ 0 and *ψ*_1_ the eigenmode with the longest spatial wavelength (i.e., a spatially uniform or constant eigenmode).

**Fig. 1.**
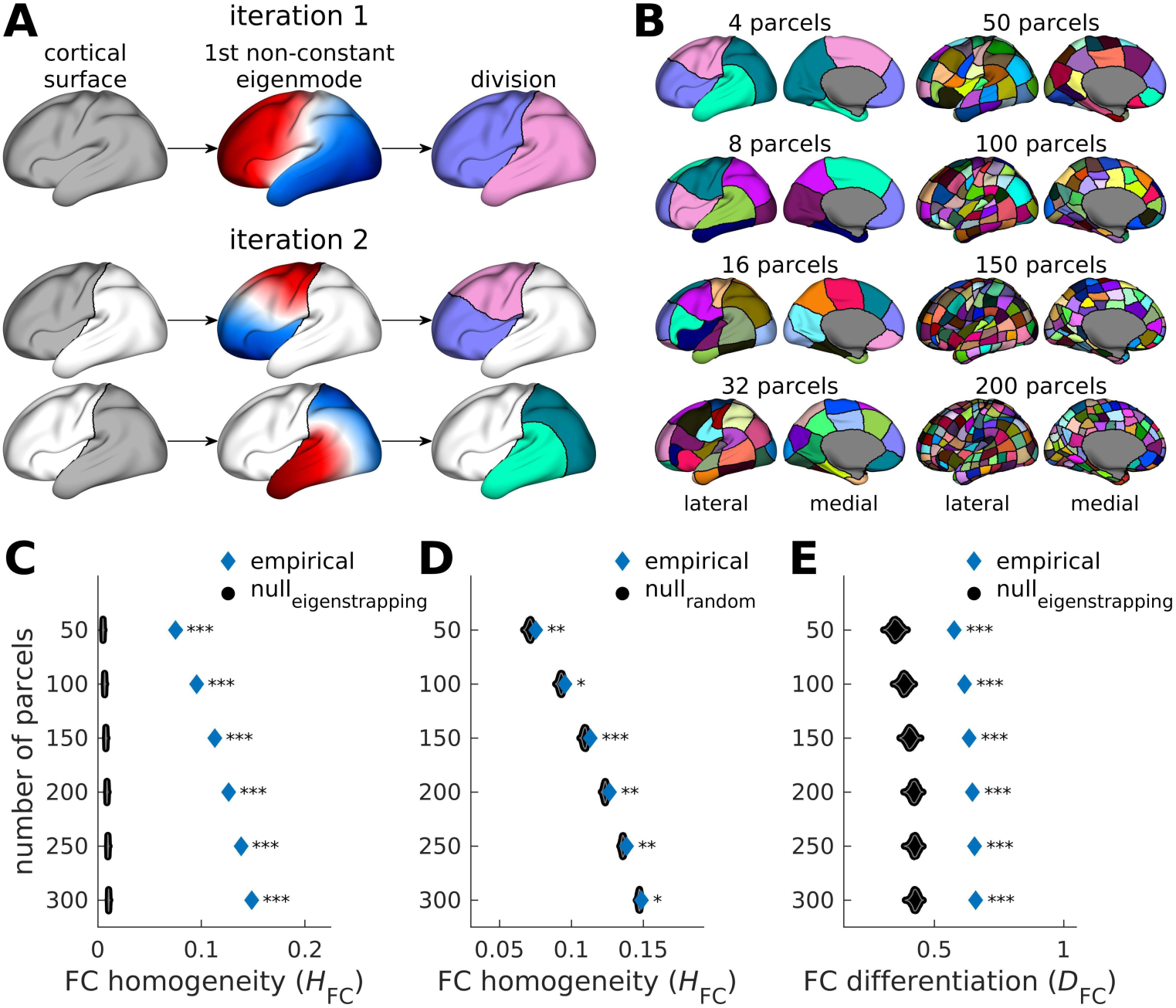
Geometrically constrained hierarchical framework for regional organization. (**A**) Schematic diagram of our iterative, hierarchical framework applied to the left human neocortex. In the first iteration, we divide the cortex into two subregions according to the nodal line of the first non-constant geometric eigenmode (white area, middle panel). In the second iteration, we repeat the process, this time by taking the first non-constant eigenmode estimated separately for the two subregions defined in the first iteration. This process is iterated several times to achieve a desired parcellation resolution, separately for the left and right hemispheres. (**B**) Example geometric parcellations across scales with 4, 8, 16, 32, 50, 100, 150, and 200 parcels. (**C**) Empirical and null FC homogeneity (*H*_FC_) of the geometric parcellations across scales (50, 100, 150, 200, 250, and 300 parcels per hemisphere) based on 1000 randomized FCs. (**D**) Same as panel C but the null *H*_FC_ is based on 1000 random parcellations. (**E**) Same as panel C but for local FC differentiation (*D*_FC_). For panels C–E, the blue diamonds correspond to the empirical data and the black dots correspond to the null data. Each geometric parcellation showed significantly higher empirical *H*_FC_ and *D*_FC_ than the null data with varying levels of significance (***: two-sided p-value <0.001; **: two-sided p-value <0.01; *: two-sided p-value <0.05).

The nodal lines of the non-constant eigenmodes (i.e., points with zero magnitude) partition the cortex into positive and negative domains of approximately equal sizes. We use the first non-constant eigenmode *ψ*_2_ with eigenvalue *λ*_2_ to partition the cortex into two subregions along the major rostrocaudal axis (Fig. 1A, top middle and right) as an initial approximation of regional boundaries at the coarsest resolution scale. This eigenmode defines the dominant axis of shape variation and is the least physically stable, meaning that it will be the first spatial pattern expressed in a reaction-diffusion process^78^. This physical property aligns with experimental evidence that most patterning molecules identified to date are expressed along a rostrocaudal gradient^52^.

After the initial partition of the cortex along the nodal line of *ψ*_2_ (Fig. 1A, top), we repeat the process successively on each new subregion over multiple iterations to generate a parcellation of any arbitrary scale (Fig. 1A, bottom). This hierarchical bipartition naturally yields 2*^N^* parcels, where *N* is the number of iterations (Fig. 1B left). To obtain a partition with arbitrary number of parcels (from 1 to 2*^N^*; Fig. 1B right), we rely on a heuristic that uses the eigenvalues of the resulting subregional eigenmodes to prioritize sub-divisions with the smallest eigenvalues (see Supplementary Figs 1A–B and ‘Hierarchical partitioning’ in Methods). This iterative procedure, based on recursive bipartition, mimics the developmental processes that shape numerous hierarchically organized, multiscale biological systems, such as limbs and digits, bronchial trees, and segmented body plans, in which tissue elements emerge through the successive branching or division of previous ones^77,79–83^. Such a process has also been implicated in brain development; for instance, the neural tube divides into the spinal cord and cerebral vesicles (hindbrain, midbrain, and forebrain), the vesicles into neuromeres, and the neuromeres into smaller sub-fields across several resolution scales^84,85^. Critically, tissue geometry constrains the precise way in which these bipartitions occur^86^.

We next investigate the degree to which our geometry-derived parcels capture some meaningful aspects of functional organization in the human neocortex. In human neuroimaging, a widely used quantity is the degree to which each parcel defines a homogeneous set of point-wise (i.e., vertex or voxel) measures^13,87–90^, with homogeneity defined in relation to inter-vertex functional coupling (FC) (i.e., average within-parcel FC; see ‘Calculation of parcel homogeneity’ in Methods). Here we quantify parcel-level FC homogeneity, *H*_FC_, as measured with resting-state functional magnetic resonance imaging (fMRI) from the Human Connectome Project (HCP) dataset^14^. We then benchmark the empirical results against two null models, generating ensembles of null *H*_FC_ calculated from: (1) null FC matrices derived via the eigenstrapping method, which randomizes fMRI data while preserving its spatial and temporal autocorrelation^91^ (Supplementary Figs 2A–B; see ‘Null FC matrices’ in Methods and Supplementary Information S2.1); and (2) random parcellations that subdivide the cortex into spatially contiguous parcels of approximately equal surface area (Supplementary Figs 2C–D; see ‘Random parcellations’ in Methods). The first null model assesses the sensitivity of our approach to capturing the regional organization of empirical FC as opposed to a generically spatially autocorrelated (i.e., non-biological) map; the second null model ensures that our approach performs better than when random regional boundaries are applied to the same empirical map. Figures 1C–D show that the geometric parcellations across multiple scales (with 50, 100, 150, 200, 250, and 300 parcels per hemisphere) are significantly more functionally homogeneous than both null models.

The *H*_FC_ metric is the most widely used validation parcellation criterion in human neuroimaging^13,87–90^ and follows from the idea that a good parcel should delineate a functionally homogeneous set of cortical points. Another important characteristic concerns how functionally differentiated or distinctive each parcel is from adjacent parcels. We therefore defined a metric, termed local FC differentiation, *D_FC_*, which is the normalized difference between within-parcel FC and between-neighboring-parcel FC (see ‘Calculation of parcel homogeneity’ in Methods), such that higher values indicate stronger differentiation between adjacent parcels. These two properties—intrinsic homogeneity (quantified with *H*_FC_) and extrinsic differentiation (quantified with *D_FC_*)—define the local specialization of a parcel. Figure 1E shows that, in addition to having higher intrinsic homogeneity (i.e., higher *H*_FC_), geometry-derived parcels across multiple scales are also significantly more functionally differentiated (i.e., higher *D_FC_*) relative to null data.

The findings in Fig. 1 demonstrate that our novel, recursive approach does not simply produce arbitrary subdivisions of the human neocortex, but captures biologically meaningful information about the regional organization of functional coupling. In the following sections, we test the generality of our approach more broadly by asking whether geometry-derived parcels have high internal homogeneity across hundreds of diverse anatomical, functional, cellular, and molecular properties of the human neocortex. We then show how our framework can be applied across brain structures beyond the neocortex and across different species, given only a model of brain geometry as input, before directly linking the algorithm that underlies our approach to the core elements of a hierarchical reaction-diffusion process.

### Geometry-derived parcels reflect functional, cellular, and molecular organization

Having demonstrated that our approach surpasses expectations set by random null models, we next ask how the FC homogeneity and differentiation of our geometric parcellations compare to 17 widely-used benchmark atlases constructed in diverse ways, including through the use of histological criteria (Histological), anatomical landmarks (Anatomical), multimodal imaging (Multimodal), or data-driven clustering of resting-state FC (Functional); see Fig. 2A, Supplementary Table 1, and ‘Benchmark brain parcellations’ in Methods. For each existing parcellation, we generate a corresponding geometric parcellation with the same number of parcels specific to the left and right hemispheres to ensure fair comparison, following our eigenvalue-based heuristic (see Supplementary Figs 1A–B and ‘Hierarchical partitioning’ in Methods).

**Fig. 2.**
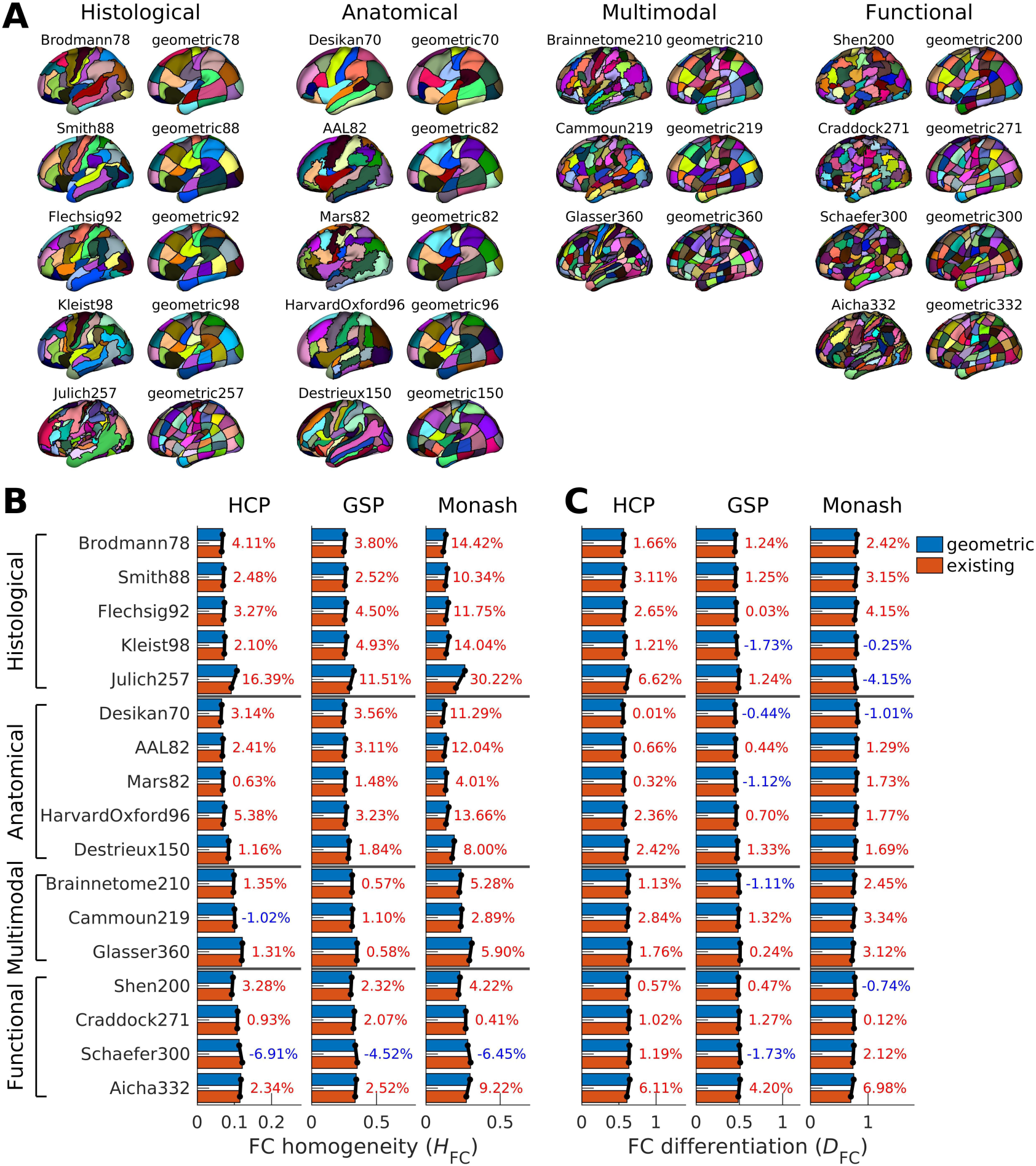
Functional homogeneity and differentiation of geometry-derived parcels of the human neocortex. (**A**) Seventeen existing benchmark parcellations and corresponding geometric parcellations with matched number of parcels. The number beside the parcellation’s name denotes the total number of parcels across both hemispheres (e.g., Brodmann78 has 78 parcels in total across the left and right hemispheres). (**B**) FC homogeneity (*H*_FC_) calculated from the HCP, GSP, and Monash datasets. (**C**) Same as panel B but for local FC differentiation (*D*_FC_). For panels B and C, the numbers represent the percentage difference between the geometric and existing parcellations, where positive/red indicates higher values for the geometric parcellations.

We first compare the FC homogeneity (i.e., *H*_FC_) of the 17 benchmark atlases and corresponding geometric parcellations using three independent resting-state fMRI datasets comprising a total of 2259 individuals [i.e., the HCP^14^; the Genomics Superstructure Project (GSP)^92^; and a dataset acquired in-house at Monash University (Monash)^93^]. For each dataset, we calculate *H*_FC_ for every benchmark atlas and its matched geometric parcellation, and express the difference as the percentage change between the two (see ‘Calculation of parcel homogeneity’ in Methods). Figure 2B shows that geometry-derived parcels have higher homogeneity than the parcels of 15 of the 17 benchmark atlases in the HCP dataset and 16 of the 17 in the GSP and Monash datasets. The Cammoun219 atlas shows higher *H_FC_* than the geometric parcellation only in the HCP dataset, but the difference is minimal (1.0%). The only benchmark parcellation with consistently and notably higher *H_FC_* is the Schaefer300 atlas, which relies on extensive fMRI training data and a sophisticated FC clustering approach^13^. In contrast, our framework uses only cortical geometry and requires no *a priori* training. We further note that the relative advantage of the Schaefer atlas is attenuated at finer spatial scales (Supplementary Fig. 3A).

We next consider the average local functional differentiation of parcels in each atlas, as quantified using *D_FC_*. Figure 2C shows that geometry-derived parcels have higher *D_FC_* than the parcels in all benchmark atlases in the HCP dataset, 12 of the 17 atlases in the GSP dataset, and 13 of the 17 atlases in the Monash dataset. Together, these findings indicate that our geometric approach defines cortical parcels that show comparable, if not superior, functional homogeneity and differentiation to most atlases currently in use today.

Figure 2A shows that, at any given scale, the geometric parcellations resemble an approximately uniform grid (Supplementary Fig. 1B) with highly uniform parcels (Supplementary Figs 1C–E) that are unlike the less regular parcel borders seen in other parcellations. This grid-like structure partly arises from the hierarchical partitioning employed by our procedure and mirrors similar rectilinear organizational motifs observed in inter-regional axonal projections^94^, which may emerge from the superposition of continuous, spatially orthogonal expression gradients of key patterning molecules^52,67^. It could also be partly driven by our application of the framework to the cortical surface geometry of a population-average template. This choice provides a common reference frame for comparison with group-level benchmark atlases, but it also smooths fine-scale anatomical variations and retains only coarse geometric features of the cortex. Applying our framework to individual cortical geometries yields boundaries that are more closely aligned with an individual’s anatomical landmarks (Supplementary Fig. 4A; Supplementary Information S3), while also yielding parcels with higher average *H*_FC_ than the average template-based approach (Supplementary Fig. 4B), particularly at coarse resolutions. This increase in homogeneity is specific to an individual’s FC data (Supplementary Fig. 4C), supporting the idea that individual differences in cortical geometry capture meaningful variation in functional organization.

Interestingly, at coarse scales (e.g., with 16 parcels), individual-specific parcellations partially align with known macroanatomical divisions, including visual and somatosensory fields (Supplementary Figs 5A–B; Supplementary Information S3). These parcels spatially overlap with corresponding areas defined in the Glasser atlas, which used multimodal imaging and manual intervention for areal definition (Supplementary Fig. 5C), while also showing higher homogeneity (Supplementary Fig. 5D). This correspondence suggests that the spatial extent of canonical areas may partly reflect the same geometric constraints captured by our framework, consistent with recent work suggesting that the spatial location of primary sensory fields may be shaped by cortical geometry^50^. Although our current goal is not to reproduce the precise boundaries of these areas, these observations suggest that one could obtain closer alignment between geometric parcels and classical areal boundaries by manually ‘locking’ selected parcels at an appropriate spatial scale before continuing the recursive partitioning procedure in other areas (Supplementary Fig. 6A), although this would come at the cost of less uniform parcel sizes and lower functional homogeneity (Supplementary Figs 6B–C). This flexibility illustrates the potential of our framework to incorporate further biological complexities, but our remaining analyses will focus on template-based cortical surfaces with regionally uniform hierarchical stopping criteria (i.e., our original and not the hybrid approach in Supplementary Fig. 6) for simplicity, and because most of the brain phenotypes we next evaluate are not individual-specific (see also Supplementary Information S3).

This reliance on population-average cortical surface template nonetheless raises the question of whether the high internal functional homogeneity of the geometry-derived parcels can be explained by low-level spatial features, such as parcel-size uniformity or compactness. To address this concern, we perform two robustness analyses. First, we find that differences in *H_FC_* between the geometric and benchmark parcellations are not significantly associated with differences in parcel-size variance (Supplementary Figs 7A–B) or parcel compactness (Supplementary Figs 7C– D). Thus, our findings cannot be explained by the more uniform appearance of the geometric parcels. Second, following the analysis in Fig. 1C, we find that the empirical *H*_FC_ of the geometric parcellations, as well as their differences relative to benchmark parcellations, exceed values expected under eigenstrapping nulls that preserve the empirical spatial and temporal autocorrelation of fMRI data (Supplementary Figs 8A–B; see Supplementary Fig. 8C for replication with an alternative null model^95^ described in Supplementary Information S2.2). Additionally, following the analysis in Fig. 1E, we find that the empirical *D*_FC_ of the geometric parcellations exceed values expected under eigenstrapping nulls (Supplementary Fig. 8D). These findings indicate that the geometric parcels are indeed capturing biologically relevant information that cannot be explained by generic autocorrelations. In other words, the functional organization captured by our framework reflects the specific orientation and placement of geometry-derived parcels, rather than generic properties of spatially contiguous, compact, and regular parcels.

Having established that geometry-derived parcels are functionally homogeneous with respect to three independent fMRI datasets, we next evaluate parcel-level homogeneity across 242 diverse non-FC cortical phenotypes or maps drawn from open-source repositories^14,96,97^ (Fig. 3A; see ‘Brain phenotypes’ in Methods). These maps include 1 morphometry map (cortical thickness); 3 microstructure maps (T1w:T2w ratio – an MRI metric that indirectly probes intracortical myeloarchitecture^98^, NDI – neurite density index, and ODI – orientation dispersion index); 50 cytoarchitecture maps (layer-specific cell density); 5 metabolism maps derived from positron emission tomography (PET) (CMRGlu – cerebral metabolic rate of glucose, CBV – cerebral blood volume, CBF – cerebral blood flow, CMRO_2_ – cerebral metabolic rate of oxygen, and synaptic density); 1 gene expression map (first principal component of gene profiles); 19 chemoarchitecture maps (i.e., neurotransmitter receptor density maps derived from PET); 47 HCP task-activation maps; and 116 meta-analytic task-activation maps from Neurosynth^97^. Following previous work^13,89^, for each phenotype, we estimate map-specific homogeneity, denoted *H*_m)*_, as the inverse of the variance of map values across the vertices within each parcel (see ‘Calculation of parcel homogeneity’ in Methods). Evaluating such a diverse set of phenotypes allows us to test whether our geometrically constrained framework captures features of regional organization that are independent of the measurement technique, rather than idiosyncrasies of any specific imaging modality (e.g., fMRI) or brain property (e.g., FC).

**Fig. 3.**
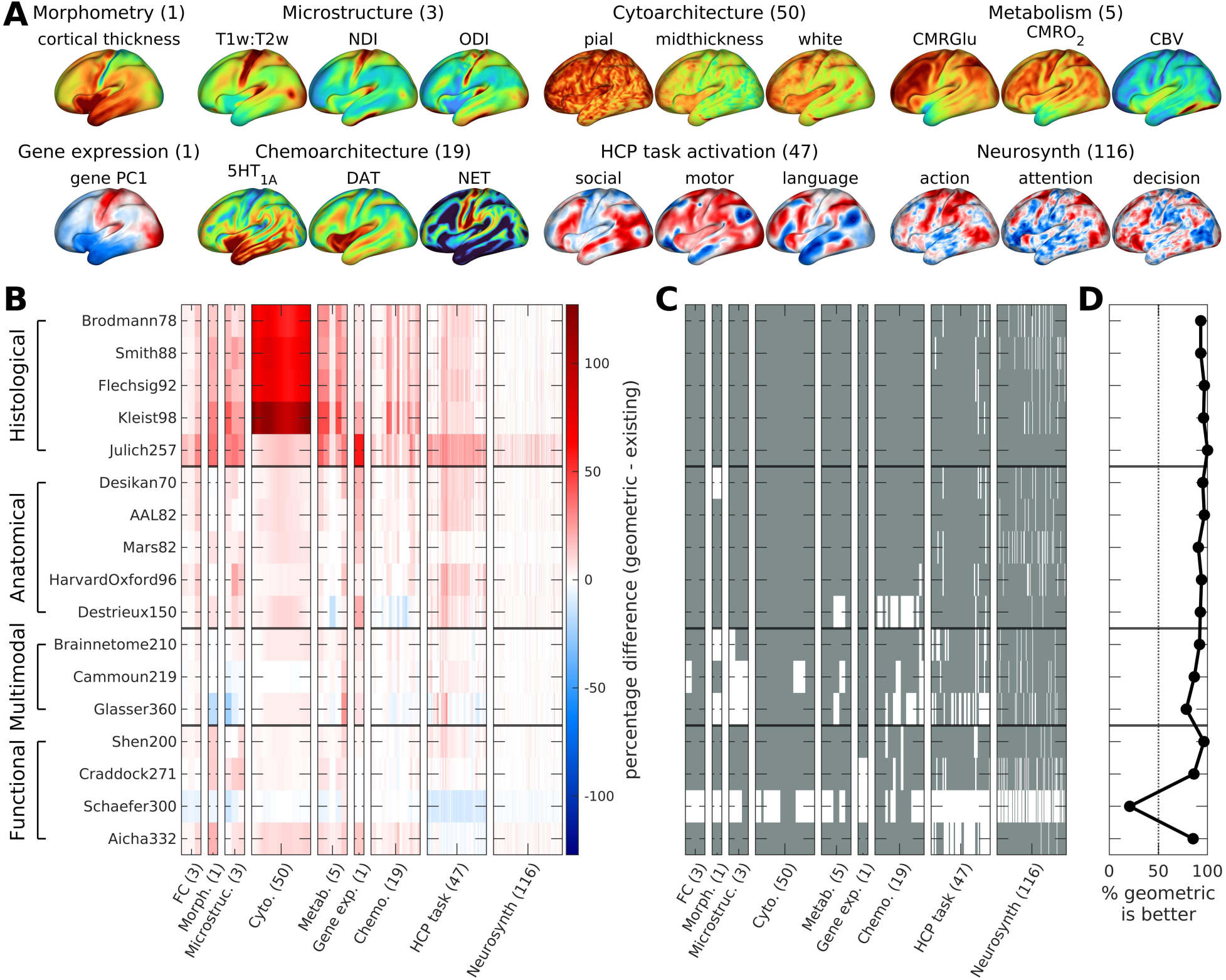
Homogeneity of geometry-derived parcels across diverse maps of the human neocortex. (**A**) Examples from 242 non-FC brain maps derived from various modalities capturing distinct phenotypes, including those related to morphometry, microstructure, cytoarchitecture, metabolism, gene expression, chemoarchitecture, HCP task activation, and Neurosynth task activations. The number beside the phenotype’s group name denotes the total number of available maps. For phenotypes with more than three maps, only three representative maps are visually shown for brevity. (**B**) Percentage difference in homogeneity between geometric and benchmark parcellations. Each row shows the result for one benchmark comparison and each column shows one brain map (including the three FC-based results in Fig. 2B), where red indicates higher homogeneity for geometric parcellations. (**C**) Binarized version of panel B, where gray indicates higher or equivalent homogeneity for geometric parcellations, with equivalence quantified as a difference threshold of ±1%. (**D**) Percentage of maps for which geometric parcellations show higher or equivalent homogeneity.

Figures 3B–C show that geometry-derived parcels have consistently high homogeneity across diverse cortical phenotypes, with some differences relative to benchmark atlases exceeding 100%. Specifically, geometric parcellations show higher or equivalent homogeneity for >78% of maps relative to 16 out of 17 benchmark parcellations (Fig. 3D; see Supplementary Fig. 9A for other equivalence thresholds). The exception is again the Schaefer300 atlas, which shows higher homogeneity than its geometric counterpart across 79.2% of the maps, highlighting the excellent generalizability of this FC-derived parcellation. However, the average difference between Schaefer300 and the corresponding geometric parcellation is modest (mean=4.5%; min=1.0%; max=14.4%), particularly when compared with the average difference between geometric and histological parcellations (mean=23.3%; min=1.0%; max=127%). We also note that the higher homogeneity of the Schaefer atlases relative to the geometric parcellations attenuates at finer spatial scales (Supplementary Figs 3B–D). The higher homogeneity of the geometric parcellation relative to the well-known Julich257 cytoarchitectonic atlas could partly reflect the latter’s very large parcels with ambiguous areal identities, termed ‘gap maps’^99^, although further analysis reveals that our results and conclusions are robust to removal of these gap maps (Supplementary Fig. 10).

Together, our findings indicate that our simple, hierarchical geometric approach is sufficient to generate a blueprint of regional organization that generalizes across multiple molecular, cellular, anatomical, metabolic, and functional phenotypes, and which shows comparable validity to existing atlases defined for more specific purposes. We further emphasize that our goal is not to identify a single ‘optimum’ atlas for all applications, since the utility of any specific atlas depends on the question under investigation (e.g., see ref.^100^ for a systematic comparison of atlases). For example, multimodal atlases, such as Glasser360, may be useful when the aim is to capture anatomical and functional information. Functional atlases, such as Schaefer300, may better capture fine-grained functional boundaries, even when those boundaries do not precisely coincide with histological or macroanatomical divisions. In contrast, our framework is modality-agnostic and uses only the geometry of the cortical surface, typically obtained from a T1w anatomical scan, making it useful for identifying a broad organizational blueprint at various scales of interest. The fact that geometry-derived parcels show high homogeneity across functional, anatomical, cellular, and molecular phenotypes suggests that geometry approximates an initial scaffold upon which boundaries may be refined by developmental, connectional, and activity-dependent processes.

### Geometric constraints on regional organization generalize to other mammals

To test whether the hierarchical geometric processes captured by our framework reflect a conserved influence of geometry on the multiscale regional organization of the mammalian brain, we investigate whether geometry-derived parcels also capture the organization of neocortical properties in the macaque, marmoset, and mouse. For the macaque and marmoset, we apply the same framework used in Fig. 1A to a triangular mesh representation of their population-averaged cortical surfaces (10,242 vertices for each hemisphere of the macaque; 37,974 and 38,094 for the left and right hemispheres of the marmoset, respectively). For the mouse, we apply the framework to its neocortex in three-dimensional (3D) volumetric space (200 μm isotropic voxel resolution; see ‘Derivation of geometric eigenmodes’ in Methods) because existing atlases of the mouse brain are only available as 3D volumes.

We compare the homogeneity of geometric parcels with 15 existing parcellations derived from histological and multimodal data (Fig. 4A; Supplementary Table 1; 10, 4, and 1 parcellations for the macaque, marmoset, and mouse, respectively). Note that some parcellations, such as Brodmann82 and Glasser360 in the macaque, were originally constructed in humans but connectivity-based alignment methods allow their robust projection on the animal cortices^101^ (see Supplementary Table 2). Homogeneity is evaluated on 114 different cortical phenotypes (20, 5, and 89 phenotypes for the macaque, marmoset, and mouse, respectively), spanning resting-state FC (measured using fMRI and calcium imaging), morphometry, microstructure, chemoarchitecture, cytoarchitecture, and gene expression (see ‘Benchmark brain parcellations’, ‘Brain phenotypes’, and ‘Calculation of parcel homogeneity’ in Methods). The use of both *in vivo* and *ex vivo* data across functional, cellular, and molecular properties allows us to test whether geometry captures organizational features that generalize across species, measurement modalities, and biological phenotypes.

**Fig. 4.**
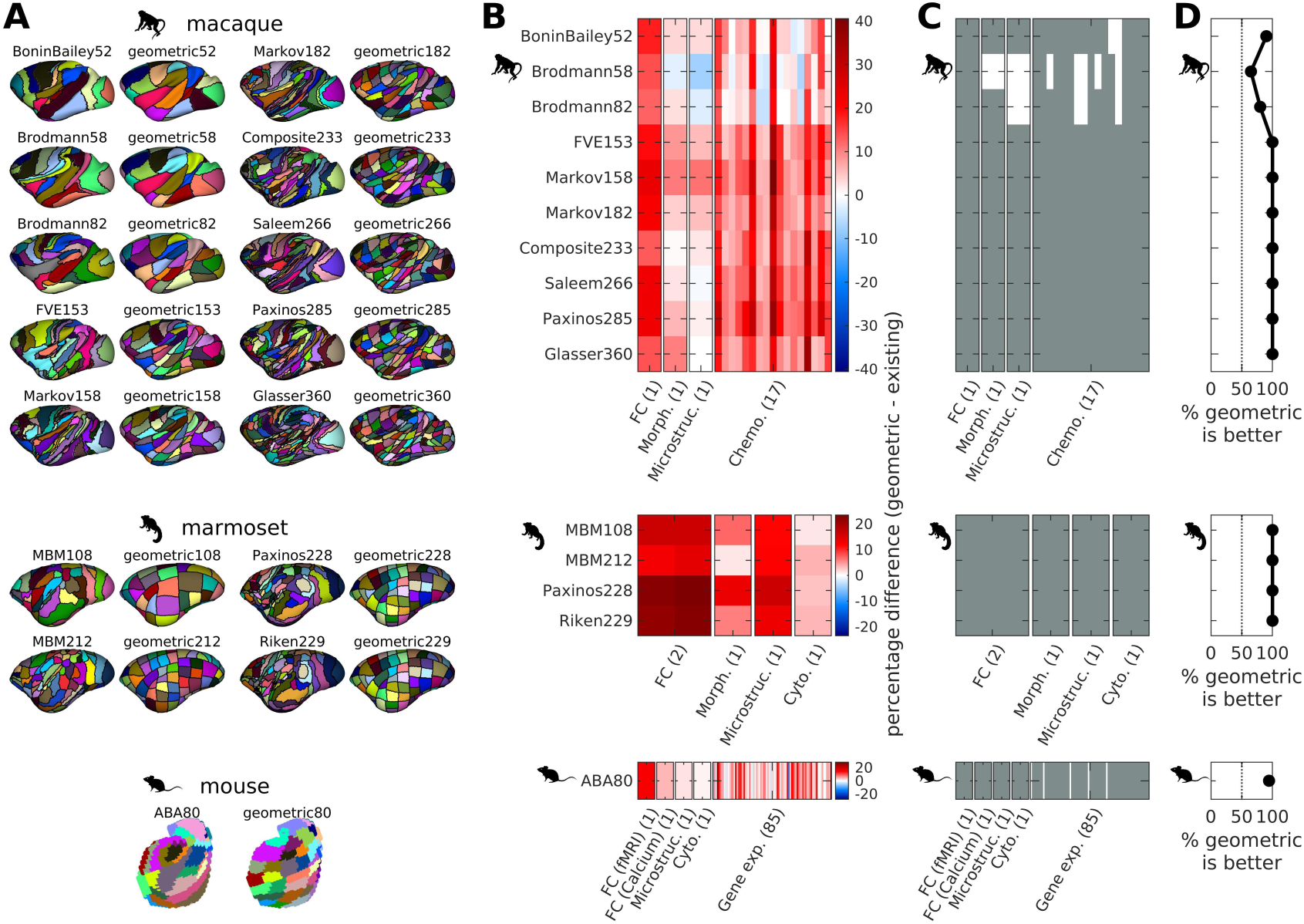
Homogeneity of geometry-derived parcels across diverse maps of the macaque, marmoset, and mouse neocortex. (A) Existing benchmark parcellations and corresponding geometric parcellations with 13 matched number of parcels. The number beside the parcellation’s name denotes the total number of parcels across both hemispheres. (B) Percentage difference in homogeneity between geometric and benchmark parcellations. Each row shows the result for one benchmark comparison and each column shows one brain map, where red indicates higher homogeneity for geometric parcellations. (C) Binarized version of panel B, where gray indicates higher or equivalent homogeneity for geometric parcellations, with equivalence quantified as a difference threshold of ±1%. For panels B and C, the number beside the phenotype’s group name denotes the total number of available maps. (D) Percentage of maps for which geometric parcellations show higher or equivalent homogeneity

Figures 4B–C show that geometry-derived parcels are consistently homogeneous across cortical phenotypes in all three species. Specifically, we find that the geometric parcellations have higher or equivalent homogeneity for >65%, 100%, and 94.4% of the brain phenotypes in the macaque, marmoset, and mouse, respectively (Fig. 4D; see Supplementary Figs 9B–D for other equivalence thresholds). In the macaque, lower homogeneity is observed only relative to the BoninBailey52, Brodmann58, and Brodmann82 parcellations in 2, 7, and 4 out of 20 maps, respectively, but the average differences are small (mean=3.5%; min=1.2%; max=7.4%). These differences are modest compared with the 12.6%, 14.3%, and 18.0% higher FC homogeneity of the geometric parcellations relative to the BoninBailey52, Brodmann58, and Brodmann82 atlases, respectively. In the mouse, the widely used ABA80 atlas shows higher homogeneity than the geometric parcellations in only 5 out of 89 maps (all related to gene expression), with an average difference of 5.4% (min=1.8%; max=12.8%).

These results are notable given that many of the non-human primate and mouse parcellations have been defined through careful cytoarchitectonic analysis, which is often considered the gold standard in the field. As in the human analyses, however, the results suggest that different parcellations capture different aspects of cortical organization. Cytoarchitectonic parcellations, such as BoninBailey52, describe an important aspect of areal differentiation especially in primary sensory cortices, whereas the geometric parcellations capture broad spatial structure that generalizes across multiple phenotypes. The consistency of results across species supports the idea that geometry provides a universally conserved scaffold for multiscale regional organization.

A key advantage of our framework is that it is simple, fast, and flexible, requiring only a geometric model of a given brain structure, which can easily be obtained from T1w MRI or microscopy. This simplicity contrasts with classical approaches that require expensive and time-consuming manual histological characterization or the application of sophisticated clustering algorithms to high-dimensional functional and physiological data. Our framework can therefore provide an initial map of geometrically constrained multiscale regional organization for species in which such organization remains poorly characterized.

To demonstrate this flexibility, we apply our framework to an open-source repository of cortical surfaces from 24 distinct species in the Euarchontoglires superorder of mammals (also known as Supraprimates), which includes the groups of Primata, Scandentia, Dermoptera, Rodentia, and Lagomorpha^102^. Figure 5 shows example geometric parcellations at arbitrary resolutions within the spatial limits of MRI (the example panels show 8, 16, and 32 parcels). These geometric parcellations demonstrate the immediate practical and neuroscientific utility of our framework for under-studied mammalian species, opening new opportunities to study the regional properties of the brains of these species that would not otherwise be possible. We provide this library of atlases as an open resource to facilitate future comparative research and validation (see ‘Data availability’ section), but we caution that our geometry-derived parcels are not assumed to be homologous across species. Establishing homology requires additional anatomical, structural, and functional characterization; for example, using the methods developed in ref.^103^.

**Fig. 5.**
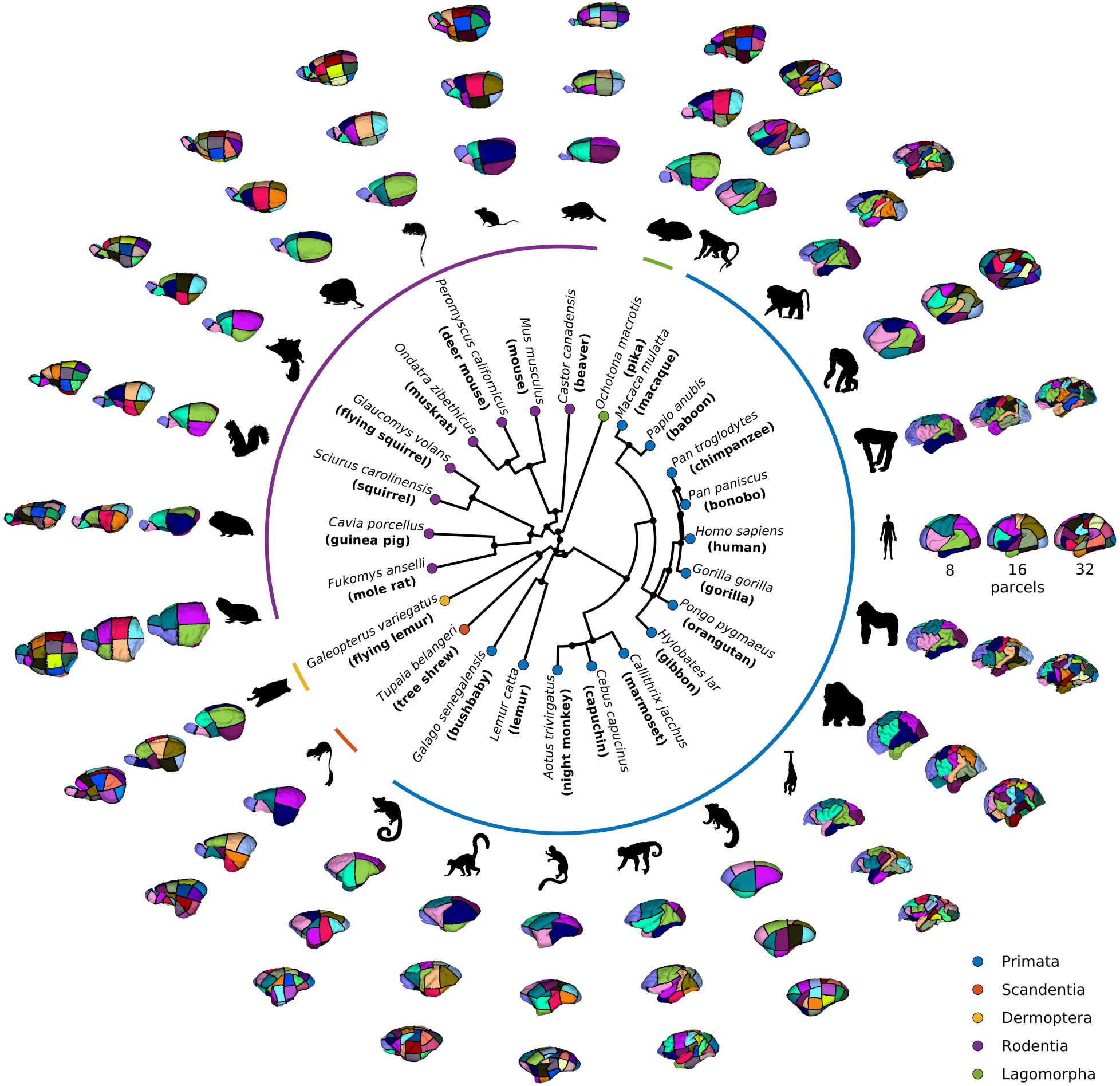
Geometry-derived multiscale organization across 24 diverse mammalian species. The center graph shows the phylogenetic relationships between the species from 5 groups of the Euarchontoglires mammalian superorder^102^; i.e., Primata, Scandentia, Dermoptera, Rodentia, and Lagomorpha. For each species, the insets show example geometric parcellations with 8, 16, and 32 parcels (from inner to outer ring).

### Geometric constraints on regional organization generalize to non-neocortical structures

Having shown that our geometric framework captures regional organization in the neocortex across multiple species, we next ask whether the same framework also applies to non-neocortical (i.e., subcortical and allocortical) structures of the human brain. We focus on the following 7 structures: hippocampus (HIP); amygdala (AMY); thalamus (THA); nucleus accumbens (NAc); globus pallidus (GP); putamen (PUT); and caudate (CAU). For each structure, we generate geometric parcellations using 3D volumetric (voxel-based) models of structure-specific geometry derived from T1w MRI (at 2 mm isotropic voxel resolution)^47^. We then compare their FC homogeneity with that of 20 existing benchmark parcellations across the 7 structures (Fig. 6A; Supplementary Table 2; specifically 4, 2, 6, 2, 2, 2, and 2 parcellations for HIP, AMY, THA, NAc, GP, PUT, and CAU, respectively) in the three independent resting-state fMRI FC datasets analyzed in Fig. 2 (see ‘Calculation of parcel homogeneity’ in Methods). These benchmark parcellations were derived using a variety of approaches based on cytoarchitecture, myeloarchitecture, resting-state fMRI, and diffusion MRI (Supplementary Table 2).

**Fig. 6.**
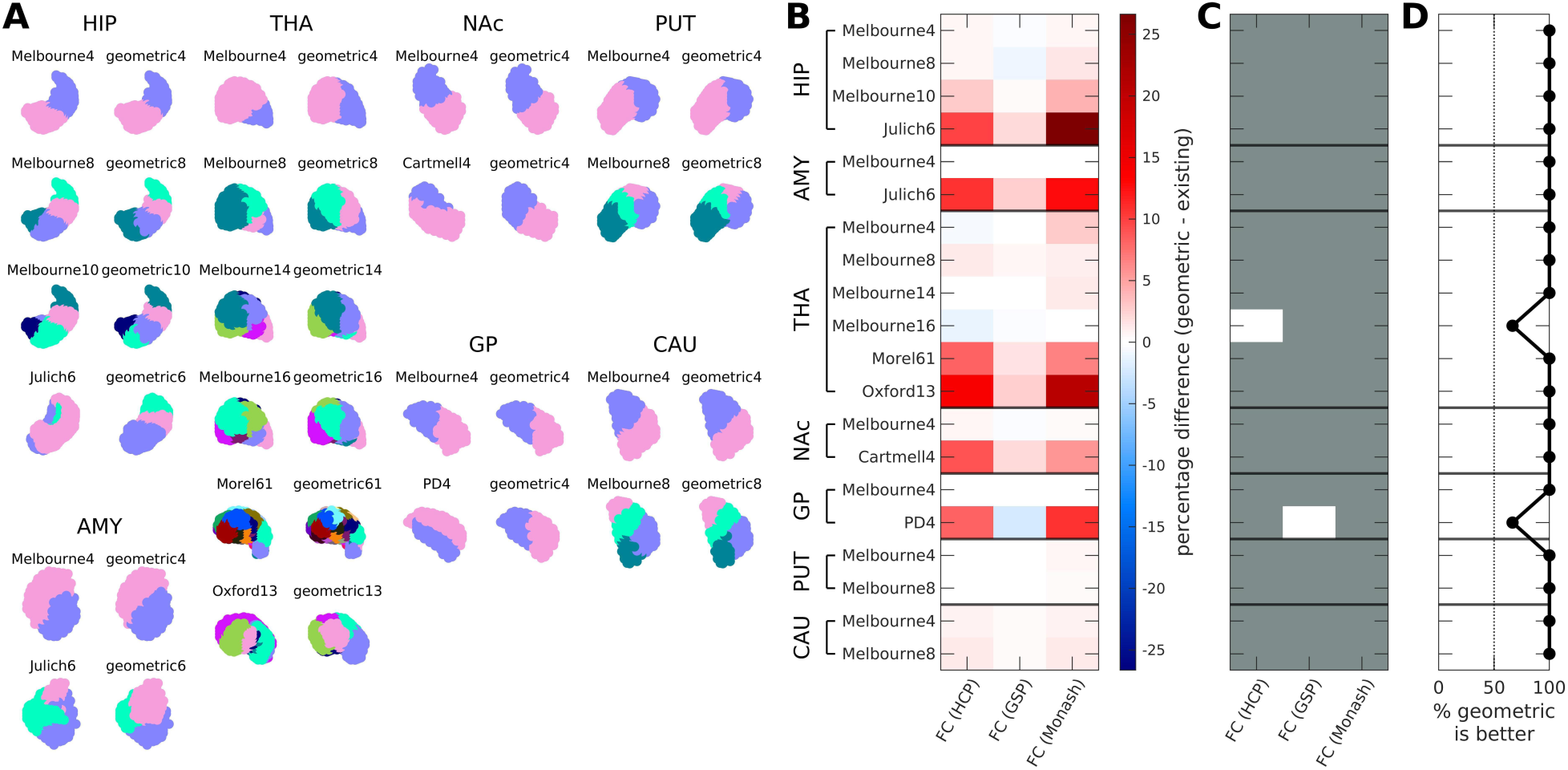
Functional homogeneity of geometry-derived parcels of 7 human non-neocortical structures. (A) Existing benchmark parcellations and corresponding geometric parcellations with matched number of parcels across the 7 structures (HIP: hippocampus; AMY: amygdala; THA: thalamus; NAc: nucleus accumbens; GP: globus pallidus; PUT: putamen; CAU: caudate). The number beside the parcellation’s name denotes the total number of parcels across both hemispheres. (B) Percentage difference in homogeneity between geometric and benchmark parcellations. Each row shows the result for one benchmark comparison and each column shows one FC dataset, where red indicates higher homogeneity for geometric parcellations. (C) Binarized version of panel B, where gray indicates higher or equivalent homogeneity for geometric parcellations, with equivalence quantified as a difference threshold of ±1%. (D) Percentage of maps for which geometric parcellations show higher or equivalent homogeneity.

Figures 6B–D show that the geometric parcellations of the 7 human non-neocortical structures generally have higher or equivalent *H_FC_* relative to benchmark parcellations. The only exceptions are the Melbourne16 THA parcellation in the HCP dataset and the PD4 GP parcellation in the GSP dataset, but the homogeneity differences are small (1.1% and 2.1%, respectively) and only slightly above our 1% equivalence threshold (see Supplementary Fig. 9E for other thresholds). These exceptions could be due to differences in fMRI processing steps and scanner parameters across datasets. These exceptions are isolated, with the geometric parcellations showing higher or equivalent homogeneity in 18 of 20 benchmark comparisons across multiple datasets.

The geometric parcellations also show strong spatial correspondence to the boundaries defined by many of the existing parcellations across scales, particularly the Melbourne4 parcellations for HIP, AMY, NAc, GP, PUT, and CAU (average Dice coefficients of 0.93, 0.94, 0.95, 0.97, 0.93, and 0.94, respectively). This correspondence is expected because the Melbourne parcellations draw regional boundaries using sharp transitions in spatially varying FC gradients, which correspond almost precisely with the geometric eigenmodes of these non-neocortical structures^47^. Thus, in these structures, geometry appears to capture a coarse organizational scaffold that is also reflected in FC gradient-based parcellations. Together, these findings further underscore that geometric constraints on regional organization are not limited to the neocortex, but extend to diverse brain structures with distinct geometries, anatomical properties, and functional roles.

### Recursive geometric partitioning approximates a sequential reaction-diffusion patterning mechanism

Our geometric framework rests on an implicit generative model of the brain’s regional organization that emphasizes: (1) the fundamental physical constraints imposed by geometry on the diffusion of early patterning molecules; and (2) recursive bipartition to generate a hierarchical, multiscale architecture. Here, we demonstrate how these two mechanisms capture the essence of a generic reaction-diffusion process hypothesized to contribute to the establishment of multiscale regional organization in the developing brain.

For brevity, we focus on the human neocortex and simulate a linearized reaction-diffusion (RD) model of an activator-inhibitor type^77^ (see ‘Reaction-diffusion model’ in Methods). We emphasize that this model is not intended to precisely capture the action of any specific molecule or interactions between a set of molecules, whose biophysical details are often unknown (for exceptions, see refs^104,105^). Instead, the model identifies a minimally sufficient set of physiological mechanisms that can yield the regional patterning captured by our framework.

We first focus on molecules seeded at maximally distant points near the rostral and caudal poles of the left hemisphere (Fig. 7A), following evidence that several morphogens and transcription factors (e.g., *Emx2* and *Pax6*) show opposing concentration gradients along the rostrocaudal (RC) axis^65,71^. Depending on model parameter combinations, Turing showed that this type of model can lead to the spontaneous emergence of spatially non-constant solutions through a spatial Turing instability^56^ (see ‘Reaction-diffusion model’ in Methods for the instability conditions). Figure 7B (top row) shows the equilibrium RD solution for (*u* − *v*) using the seeds in Fig. 7A and parameters that induce a Turing instability (see ‘Reaction-diffusion model’ in Methods for the specific parameters). The RD model solution gives rise to an increase of *u* and *v* in opposite halves of the cortex, with *u* > *v* towards the rostral pole and *v* > *u* towards the caudal pole. This pattern divides the cortex into two subregions separated by a nodal line that bears a striking qualitative resemblance to the RC geometric eigenmode (Fig. 7B bottom row), which is the starting point of our framework (Fig. 1A). The correspondence between the RD solution and RC eigenmode is high, both for the continuous maps, as quantified by spatial correlation (i.e., 0.85), and for their binary subdivisions, as quantified by Dice coefficient (i.e., 0.82). The spatial pattern of the model RD solution in Fig. 7B is robust to replacing the linear activator-inhibitor interactions in the model with nonlinear functions (Supplementary Fig. 11).

**Fig. 7.**
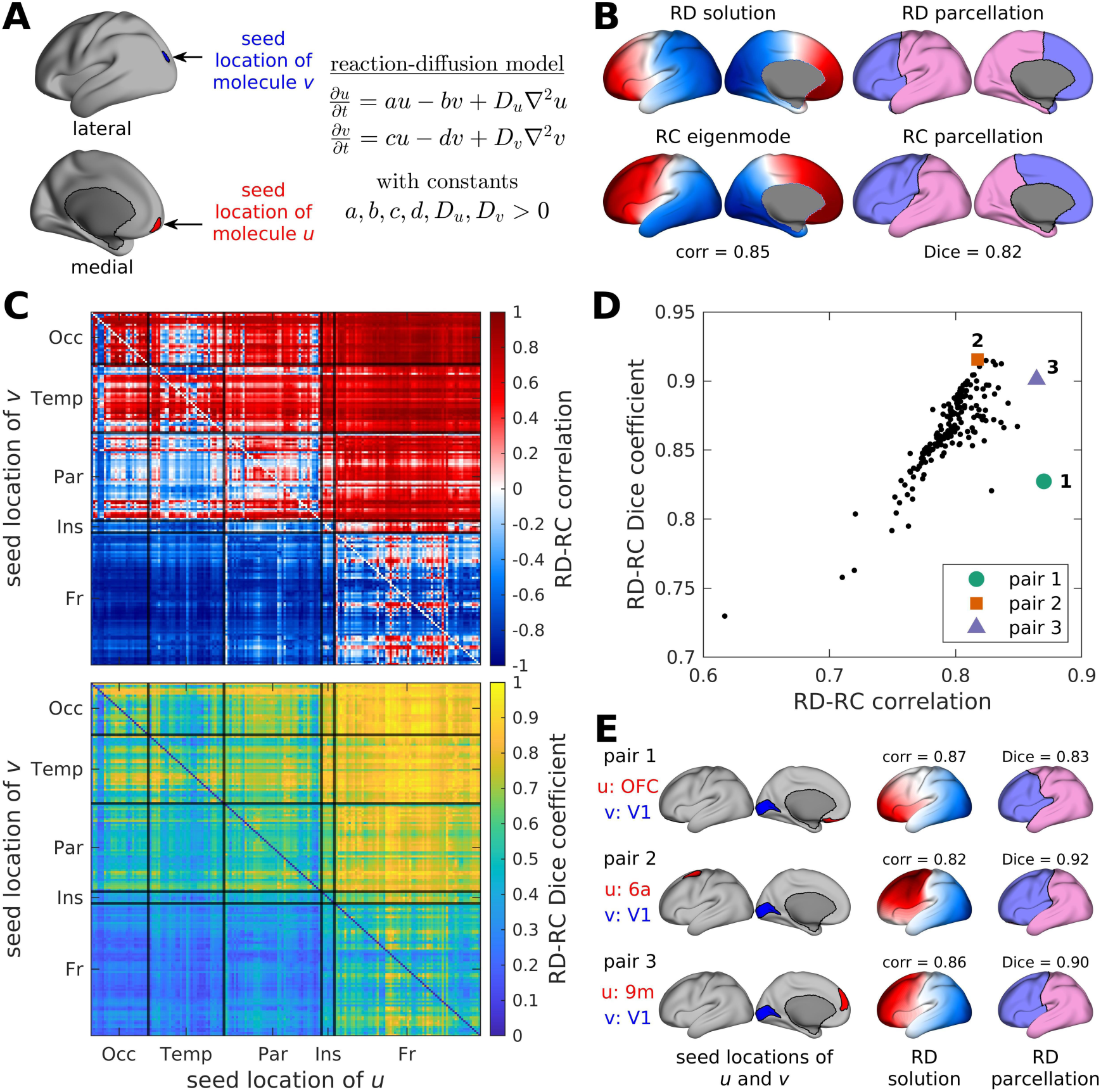
**Linear reaction-diffusion model of regional organization**. (A) We use a reaction-diffusion (RD) model that assumes two molecules, *u* and *v*, are seeded from distinct locations and diffuse in space and time with diffusion constants *D_u_* and *D_v_*, respectively. The schematic shows example seed locations of the molecules at maximally distant points near the rostral and caudal poles of the left hemisphere of the human neocortex, specifically at regions 10pp (orbital prefrontal cortex) and V3B for *u* and *v*, respectively, based on the Glasser parcellation^2^. The constants *a*, *b*, *c*, and *d* are assumed to be positive, such that *u* acts as the activator and *v* as the inhibitor. (B) The top row shows an example model RD solution (*u* − *v*) and the corresponding binary subdivision using parameter combinations that induce a Turing spatial instability. The instability conditions are: *D_u_* < *D_v_*; *a* < *d*; *ad* < *bc*; and *d*⁄*D_v_* < *a*⁄*D_u_*. The molecules are seeded at the locations described in panel A. The bottom row shows the rostrocaudal (RC) geometric eigenmode and the corresponding binary subdivision. Correspondence between the model RD solution and RC eigenmode is quantified using spatial correlation, and correspondence between the binary subdivisions is quantified using Dice coefficient. (C) Spatial correlations between model RD solutions and the RC geometric eigenmode (Top), and Dice coefficient between their binary subdivisions (Bottom), when *u* and *v* are seeded in different locations. Each location is assigned to one of five anatomical cortices: Occipital (Occ); Temporal (Temp); Parietal (Par); Insular (Ins); and Frontal (Fr). Correspondence remains high when the activator *u* is seeded anywhere in the anterior part of the brain and the inhibitor *v* is seeded anywhere in the posterior part of the brain. (D) RD-RC spatial correlation and Dice coefficient when the inhibitor *v* is seeded at V1 and the activator *u* is seeded in different locations. Each point represents a different seed location for *u*. Three seed-location pairs are highlighted that show high spatial 18 correlation, high Dice coefficient, or both. (E) Seed locations, resulting model RD solution, and resulting binary subdivision (from left to right) for the three highlighted pairs in panel D.

Figure 7C shows the effect of seeding molecules *u* and *v* at different locations using the parameter combinations that produce the RD solution in Fig. 7B. The correspondence between the model RD solution and RC geometric eigenmode, and their resulting binary subdivisions, is extremely robust, with high spatial correlations (mean=0.75; median=0.79; min=−0.07; max=0.99) and high Dice coefficients (mean=0.84; median=0.85; min=0.48; max=0.98) obtained as long as the activator *u* is seeded anywhere in the frontal cortex and the inhibitor *v* is seeded anywhere in the occipital or temporal cortices.

Some of the strongest correspondences are obtained when the inhibitor *v* is seeded at region V1 (primary visual cortex) regardless of the seed location of the activator *u*, with mean spatial correlation of 0.80 (min=0.62; max=0.87) and mean Dice coefficient of 0.86 (min=0.73; max=0.92), as shown in Fig. 7D. This result is notable given that V1 is the most cytoarchitectonically and transcriptomically distinct region of the primate neocortex^60,106,107^ and is the last such region to complete neurogenesis^108^. Figure 7E shows specific optimal solutions with either high spatial correlation, high Dice coefficient, or both when the activator *u* is seeded in the OFC (orbitofrontal cortex), area 6a (premotor cortex), and area 9m (anterior part of the dorsomedial prefrontal cortex), respectively. This robustness supports a consistent geometrically constrained blueprint for regional patterning in the face of biological noise^109^.

In principle, the RD model can be iterated to generate recursive divisions within each initial partition, yielding a multiscale regional organization that resembles the one produced by our geometric algorithm. The model thus implies that spatial variations in molecular concentration defined by the equilibrium RD solution act as thresholds that trigger the diffusion of additional molecules and more fine-grained cellular or molecular processes within each subregion. This process echoes the threshold-dependent molecular dynamics proposed in Wolpert’s classical model of the French Flag problem, which aims to understand how positional information is conferred upon developing cellular arrays^110,111^. Our model implies that the new molecule distributions further destabilize along the dominant geometric eigenmode of each subregion, yielding further bipartitions. In this way, the recursive nature of our framework can be interpreted as an idealized hierarchical sequence of Turing instabilities, with each stage triggering the next set of reaction-diffusion processes in a way that is robust to the specific parameterization and details of the model (see Supplementary Information S1).

This hierarchical sequence is efficient and biologically plausible. Once concentration levels saturate following the first division, a second instability is triggered within each subregion, yielding further two-fold subdivisions. Under this process, only log_2_ *N* hierarchical levels are required to produce *N* regions (see Supplementary Information S1.4). A similar mechanism of threshold-dependent cascading is thought to underlie the progressive segmentation of the *Drosophila* body plan along the RC axis^112,113^. This recursive process may thus capture a conserved aspect of regional patterning that supports the progressive compartmentalization of different structures^82,85,114^ and which can be effectively approximated by the first non-constant geometric eigenmode (i.e., the RC eigenmode at the first division). Accordingly, this eigenmode strongly aligns with a highly conserved gradient of neurogenic timing and neuronal density in mammals^115,116^.

## Conclusions

Understanding the regional organization of the mammalian brain is essential for explaining how specialized functions emerge and how they are integrated to support coordinated behavior^4,5^. Distinct brain regions have traditionally been defined using descriptive approaches that identify areas or parcels from expert anatomical criteria or statistical clustering of brain data. Although these approaches have been foundational, they do not directly explain why regional organization takes the form that it does. Here, we have developed a geometrically constrained framework for characterizing the hierarchical, multiscale organization of mammalian brain structures, allowing us to investigate the degree to which geometric constraints can generate a generalizable blueprint of regional organization in the brain.

We showed that geometry-derived parcels satisfy two defining characteristics of a valid and useful parcellation^5,117^; i.e., the resulting parcels show very high internal homogeneity across diverse molecular, cellular, anatomical, metabolic, and functional properties (Figs 1–4 and 6), while also showing a high degree of functional differentiation relative to neighboring parcels (Figs 1E and 2C). In most cases, the internal homogeneity and functional differentiation of our geometric atlases matched or surpassed existing benchmark atlases. This observation, when coupled with our extensive tests against multiple null models, underscores that our framework captures biologically meaningful properties of local specialization, even if it defines boundaries that differ from standard atlases.

We nonetheless emphasize that our goal here is not to define a single ‘optimum’ parcellation, since the utility of an atlas depends on the question of interest; e.g., cytoarchitectonic parcellations are useful when classical anatomical nomenclature is required, while functional parcellations are useful for reducing the dimensionality of fMRI data. Instead, we use our framework to show how geometric constraints are sufficient to generate a fundamental, modality-agnostic blueprint of regional organization that generalizes across biological phenotypes, brain structures, and species. The simplicity of our framework, which only requires a model of brain geometry as input (e.g., from T1w MRI), offers additional practical advantages for investigating species or structures for which multiscale regional organization remains poorly characterized. Although geometry-derived parcels do not necessarily align with existing areal nomenclature and should not be assumed to be homologous across species, their high internal homogeneity across multiple properties suggests that they are meaningful units for further functional or behavioral contextualization, much like the functionally defined Schaefer atlas^13^, which delineates parcels that often diverge from classical areal boundaries but which our analysis nonetheless indicates shows excellent generalizability across many properties, consistent with its utility and widespread usage in neuroimaging analyses.

We further showed that our framework approximates a sequential reaction-diffusion process that represents an idealization of a conserved developmental mechanism for establishing a multiscale blueprint of regional organization in the mammalian brain. We stress that both the reaction-diffusion model and the hierarchical framework are deliberate simplifications. Reaction-diffusion mechanisms have broad biological applicability^104,105,118^, but areal patterning can arise through processes that do not need two interacting molecules^119,120^, and patterning gradients that predominantly run along the non-rostrocaudal axes of the brain (at the first division) have also been identified^52^. Our model is also agnostic to the specific molecules involved in the patterning. Future work could extend the framework to incorporate known patterning molecules, thalamocortical afferents, synaptic competition, and activity-dependent plasticity, all of which are likely to refine the initial geometry-constrained scaffold and shape more specific areal boundaries and laminar organization^52,53,69,121^.

In summary, our findings indicate that simple geometric constraints are sufficient to generate a blueprint of multiscale regional organization of the mammalian brain that is conserved across species and generalizes across distinct brain structures and many cellular, molecular, anatomical, metabolic, and functional properties. This blueprint may act as a scaffold upon which more specific anatomical, molecular, and activity-dependent processes act to further shape local functional specialization. Our approach offers a unified framework for studying how physical constraints contribute to multiscale regional organization across brain structures, measurement modalities, and mammalian species.

## Methods

### Geometry of neocortical and non-neocortical structures

The only input to our geometrically constrained framework is a T1w magnetic resonance imaging (MRI)-derived representation of a brain structure’s geometry either at the vertex level (for surfaces) or voxel level (for volumes). For the human neocortex, we used the fsaverage template of its midthickness surface^122^, resampled in fsLR_32k space with 32,492 vertices for each hemisphere. For the macaque neocortex, we used the Yerkes19 template of its midthickness surface^123^, resampled in fsLR_10k space with 10,242 vertices for each hemisphere. For the marmoset neocortex, we used the Marmoset Brain Mapping v3 (MBMv3) template of its midthickness surface^124^, resampled in fsaverage_38k space with 37,974 and 38,094 vertices for the left and right hemispheres, respectively. For the mouse neocortex, we used the Allen Mouse Brain Common Coordinate Framework v3 (CCFv3) template in three-dimensional (3D) volumetric space at 200 μm isotropic voxel resolution.

For the 24 Euarchontoglires mammalian species, we used reconstructions of their cortical surfaces in fsaverage6 space with 40,962 vertices for each hemisphere^102^. Phylogenetically, the 24 species belong to 5 groups; i.e., Primata, Scandentia, Dermoptera, Rodentia, and Lagomorpha. The Primata group comprises 13 species: *Macaca mulatta* (macaque); *Papio anubis* (baboon), *Pan troglodytes* (chimpanzee); *Pan paniscus* (bonobo); *Homo sapiens* (human); *Gorilla gorilla* (gorilla); *Pongo pygmaeus* (orangutan); *Hylobates lar* (gibbon); *Callithrix jacchus* (marmoset); *Cebus cappuchinus* (capuchin); *Aotus trivirgatus* (night monkey); *Lemur catta* (lemur); and *Galago senegalensis* (bushbaby). The Scandentia group comprises 1 species: *Tupaia belangeri* (tree shrew). The Dermoptera group comprises 1 species: *Galeopterus variegatus* (flying lemur). The Rodentia group comprises 8 species: *Fukomys anselli* (mole rat); *Cavia porcellus* (guinea pig); *Sciurus carolinensis* (squirrel); *Glaucomys volans* (flying squirrel); *Ondata zibethicus* (muskrat); *Peromyscus californicus* (deer mouse); *Mus musculus* (mouse); and *Castor canadensis* (beaver). The Lagomorpha group comprises 1 species: *Ochotona macrotis* (pika).

For the human non-neocortical structures, we converted the existing parcellations of the 7 structures, i.e., hippocampus (HIP), amygdala (AMY), thalamus (THA), nucleus accumbens (NAc), globus pallidus (GP), putamen (PUT), and caudate (CAU), into volumetric binary masks in the Montreal Neurological Institute (MNI) space at 2 mm isotropic voxel resolution. This is to ensure that the coverage and number of voxels in the geometric parcellations match those of the existing parcellations for fair comparison. Further details of the existing non-neocortical parcellations are described in the ‘Benchmark brain parcellations’ section.

### Derivation of geometric eigenmodes

Our framework uses the geometric eigenmodes of the brain to define geometry-derived partitions and parcels. The eigenmodes are obtained by solving the Helmholtz equation defined in Eq. (1) of the main text. The Laplace-Beltrami operator (LBO) in Eq. (1) captures the intrinsic geometry of the brain structure of interest (e.g., cortical surface), i.e., geometric and spatial relations between mesh vertices^125^, and is defined generally as^126^,

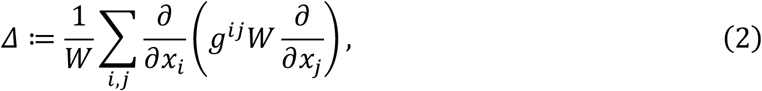

where *x_i_*, *x*_j_ are the local coordinates, (*g^i^*^j^) ≔ *G*^−1^ with *G* being the matrix of inner product of metric tensors 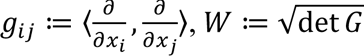, and det denotes the determinant. For surfaces, such as the human, macaque, and marmoset neocortices, we constructed the LBO using a triangular mesh representation of the T1w MRI-derived cortical sheet. For solid 3D structures, such as the mouse neocortex and human non-neocortical structures, we converted their T1w MRI-derived volumetric binary masks into tetrahedral meshes to account for the full 3D geometry of the structures (see ref.^47^ for further details). Note, however, that the LBO can also be constructed using a model of a brain structure’s geometry derived from alternative methods (e.g., microscopy) and is not restricted to T1w MRI.

To numerically calculate the geometric eigenmodes, we used our previously published code^47^ based on the LaPy Python library^125,127^ installed on the MASSIVE High Performance Computing facility^128^. The eigenvalue solutions of the Helmholtz equation in Eq. (1) are ordered sequentially according to the spatial wavelength of the spatial patterns of each eigenmode; i.e., 0 ≤ *λ*_1_ ≤ *λ*_2_ ≤⋯. Note that the first eigenvalue *λ*_1_ is approximately equal to zero (wavelength ≫ size of the brain) and the corresponding eigenmode *ψ*_1_ is a constant function with no nodal lines (zero sets of the function). Low-order eigenmodes 2, 3, and 4 correspond to spatial patterns with one nodal line and have antinodes at the opposing ends of the rostrocaudal, dorsoventral, and mediolateral axes, respectively. We further note that while our work is inspired by processes governing the developing brain, we only implemented our method on the adult brain because low-order eigenmodes of both the developing and adult brain are largely invariant^48,49^ (Supplementary Fig. 12), supporting our use of adult geometry as a reasonable proxy. Future work may consider the impact of cortical growth on the mechanisms that we model here^129,130^, although we caution that accounting for such factors can come at the cost of significant increases in model complexity. Moreover, transcriptional programs that pattern the cortex do not resemble adult expression patterns until after birth, by which point the essential features of adult cortical geometry are already established^131^. The strong performance of our simple framework suggests that adult geometry can capture a substantial component of the spatial constraints relevant to regional organization. Future work can quantify the added value of modeling developmental growth explicitly.

Additionally, our framework estimates the geometric eigenmodes while treating neocortical and non-neocortical structures as homogeneous entities, ignoring well-characterized spatial variations in cell density, myeloarchitecture, chemoarchitecture, and others^15,17,96^ and their effects on eigenmodes^132^. We used this approximation to employ the simplest possible model, and to align with all past studies of geometric eigenmodes^47,49,74,133^. It is possible that accounting for such spatial heterogeneities may perturb the structure of the eigenmodes, but incorporating them comes at the cost of an increase in model complexity and additional free parameters. The performance of our framework suggests that the added complexity may add minimal value. Future work could explore the utility of deriving eigenmodes of spatially heterogeneous neocortical and non-neocortical structures.

### Hierarchical partitioning

Our geometrically constrained framework uses the nodal lines of the eigenmodes to partition a brain structure into subregions. Specifically, we use only the first non-constant eigenmode *ψ*_2_, which is the most dominant axis of variation along the rostrocaudal direction at the first division, to partition the structure into two subregions of approximately equal sizes. Although models of regional patterning based on gradients aligned with other axes have been proposed^44–46^, we focus only on the first non-constant geometric eigenmode based on physical principles related to Turing instabilities to identify the simplest possible mechanism through which a geometry-constrained regional organization may arise (see Supplementary Information S1). Other instabilities (with a shorter wavelength) are more stable and cannot occur first.

We applied the bipartitioning in an iterative, hierarchical manner, calculating the first non-constant geometric eigenmode of each subregion and using it to further subdivide the subregion. By construction, performing *N* iterations will yield a parcellation with 2*^N^* parcels. However, the multiscale organization of the brain means that the optimal number of parcels to define remains unclear. Hence, we generalized the framework across scales, such that one can set the number of parcels to an arbitrary number between 1 and 2*^N^*. We do this by constructing a tree of eigenvalues from each iteration. We performed a maximum of 9 iterations to reduce computational burden (Supplementary Fig. 1A). We followed the tree of division from the first to *N* iterations, tracking the parent and daughter subdivisions and arranging them according to the magnitude of the eigenvalues of their first non-constant geometric eigenmodes. The lower the eigenvalue is, the more dominant that subdivision is. Hence, to generate a parcellation with *m* > 1 parcels, we retained the subdivisions corresponding to the first *m* − 1 smallest eigenvalues. To ensure that the intrinsic uniformity of the parcel sizes of the geometric parcellation (Supplementary Figs 1C–D) is maintained, we additionally constrained the method by choosing all subdivisions within an iteration first before moving to the subdivisions of the subsequent iteration. We performed this process to separately parcellate the left and right hemispheres.

The strength of our hierarchical framework is that it affords a multiscale insight into the brain’s regional organization. The resulting partitions can thus be tailored to the spatial scale relevant to a given question, unlike the multitude of brain atlases available for use today that are only defined at a single resolution scale and thus ignore the intrinsically hierarchical organization of the brain (for exceptions, see refs^13,103,134^). For instance, cytoarchitectonic studies clearly differentiate primary sensory areas, such as V1 and S1, from adjacent regions, but it is still possible to identify further functional subdivisions within these borders, such as areas processing sensory information from distinct body regions within S1 or distinct, overlapping modes of functional organization within V1^40^. The correspondence between such subregional organization and the areas defined by our framework remains to be explored, but our method nonetheless allows one to test hypotheses about how such nested functional organization may arise.

Our hierarchical bipartitioning framework mimics the processes that shape the embryonic development of various hierarchically arranged organs such as limbs, digits and vascular trees^77,79–83,135^. A bipartitioning process is favored by nature because it provides subdivisions that are robust, reliable, and insensitive to the size of the domain^82^. Hence, dividing domains of different sizes (e.g., human vs marmoset brains) into two subdomains one step at a time ensures that the subdomains will be relatively uniform in size compared to dividing the domains into several regions at once. This is important because high patterning precision is needed for robust development of brain organization (e.g., regions should generally be located in the same spatial location across individuals of a species)^109^. An interesting avenue of further work could investigate whether the regional organization of different brain areas is dictated by varying degrees of hierarchical depth in this recursive process.

### Calculation of parcel homogeneity

We compared the degree to which the geometric and benchmark parcellations capture internally homogeneous brain organization, quantified in terms of parcel homogeneity, which is the most widely used metric in the parcellation literature (see for example refs^13,87,89,90,136^). We treat high within-parcel homogeneity as evidence that a partition captures coherent spatial variation across the vertices or voxels enclosed by the parcel in the phenotype of interest. This internal coherence is a defining feature of any valid brain division and is commonly defined in relation to inter-vertex or inter-voxel functional coupling (FC)^13,87–90^ (i.e., accurate parcels are those that define sets of vertices or voxels that show high internal FC with each other), but we go beyond this common practice by additionally measuring homogeneity in terms of more diverse brain phenotypes or maps related to morphometry, microstructure, cytoarchitecture, metabolism, gene expression, chemoarchitecture, and task-related functional architecture (see ‘Brain phenotypes’ section). This is to avoid reliance on one single measure or imaging modality and to provide a comprehensive test of whether geometry captures regional organization across diverse aspects of brain structure and function.

Note that recent work has proposed a different metric to compare brain parcellations^137^, but it is most suitable only for functional data (e.g., resting-state and task-based fMRI), relies on an *a priori* choice of various heuristics (e.g., spatial bin size and weighted averaging scheme), and requires certain conditions in the data to be satisfied that are not always guaranteed (e.g., stability of within- and between-vertex correlations of functional profiles). Hence, we decided to use homogeneity (described below) to have a single, simple metric that can be generalized across diverse brain phenotypes or maps.

For each parcellation (geometric or existing), we calculated the overall parcellation homogeneity, *H*, as per prior work^13,89^, as a weighted average,

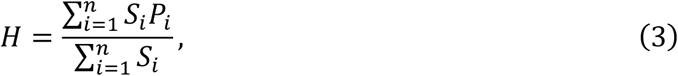

where *n* is the total number of parcels across both hemispheres, *S_i_* is the size of parcel *i* (i.e., total number of vertices or voxels enclosed by the parcel), and *P_i_* is the summary statistic for parcel *i* calculated from the phenotype or map of interest. For resting-state FC, *P_i_* is the average FC between all pairs of vertices or voxels within the parcel (i.e., within-parcel FC), such that a higher *P_i_* means that the parcel is highly homogeneous. For non-FC spatial maps, *P_i_* is the variance of map values across all vertices or voxels within the parcel, such that a lower *P_i_* means that the parcel is highly homogeneous. We used a weighted average in Eq. (3) to take into account the fact that parcels with larger sizes will tend to have lower homogeneity (Supplementary Fig. 13A), following previous work^13,89^, and to reduce the gyral and sulcal bias prevalent in cortical surface reconstructions^138,139^. The weighted average effectively decreases and increases the contribution of small and large parcels, respectively (Supplementary Fig. 13B). We pooled all *P_i_* values from both the left and right hemispheres to calculate a whole-brain homogeneity value, except for cases in which maps were only available in one hemisphere.

In addition to Eq. (3), we calculated a local FC differentiation metric, *D*_FC_, to quantify whether parcels were not only internally homogeneous but also separated from neighboring parcels. This metric thus seeks to capture a second defining feature of a brain parcel; i.e., that its functions should be internally homogeneous, while also being distinct from other parcels^5,117^. For each parcel *i*, we calculated two quantities: (1) the within-parcel homogeneity, defined as the average FC between all pairs of vertices or voxels within the parcel, *W_i_*; and (2) the between-neighboring-parcel homogeneity, defined as the average FC between vertices or voxels in parcel *i* and vertices or voxels in spatially neighboring parcels, *B_i_*. Two parcels were considered neighbors if they shared at least one boundary edge in the surface mesh or adjacent voxels in volumetric space. We then defined the normalized parcel-level FC differentiation, *D_i_*, as

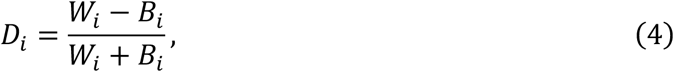

where larger values indicate stronger differentiation between within-parcel and neighboring-parcel FC. Finally, we calculated the overall parcellation differentiation, *D*_FC_, as a weighted average, analogous to Eq. (3):

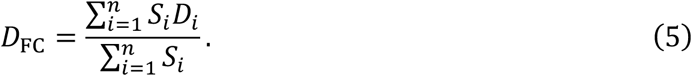

We then compared the homogeneity or differentiation of the geometric and existing parcellations (*M*^geometric^ vs *M*^existing^) as a percentage difference,

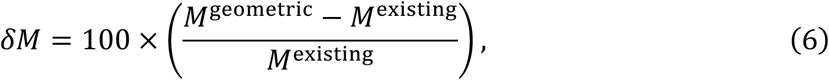

where *M* can either be *H* or *D*_FC_. For non-FC maps, we took the inverse of *H* in Eq. (3) before substituting in Eq. (6) to ensure that *δH* > 0 consistently means that the geometric parcellation has a higher homogeneity.

### Benchmark brain parcellations

We compared the homogeneity of our geometric parcellations to several existing benchmark parcellations of the human, macaque, marmoset, and mouse neocortices (Supplementary Table 1) and human non-neocortical structures (Supplementary Table 2). These were chosen because they are among the most used in the field and they were derived using diverse approaches and measures, including expert delineation and/or data-driven analysis of histological and/or imaging data (see references in the last column of Supplementary Tables 1–2). This diversity allowed us to comprehensively assess the geometric parcellations relative to other atlases. For each existing parcellation, we constructed a geometric parcellation specific to the left and right hemispheres with matched number of parcels (see the third column of Supplementary Tables 1–2) to ensure fair comparison.

Most of the parcellations were obtained directly from the references in the last column of Supplementary Tables 1–2, with some exceptions. Histological parcellations of the human neocortex (i.e., Brodmann78, Smith88, Flechsig92, and Kleist98) corresponded to the reconstructed versions in MNI space provided by ref.^140^. Parcellations of the macaque cortex (i.e., BoninBailey52, Brodmann58, FVE153, Markov158, Markov182, Composite233, and Paxinos285) were obtained from ref.^141^. All parcellations of the marmoset neocortex were obtained from the Marmoset Brain Mapping Project (https://marmosetbrainmapping.org/). All parcellations of the human non-neocortical structures, except the Melbourne parcellations, were obtained from https://www.lead-dbs.org. The Melbourne parcellations were obtained from https://github.com/yetianmed/subcortex. Finally, parcellations of the human, macaque, and marmoset neocortices originally defined in volume space were projected onto surface space using the volume-to-surface-mapping command of the Connectome Workbench software. Small parcels with fewer than 20 vertices were removed because they may have resulted from inaccuracies in the volume-to-surface projection, especially for parcellations with subcortical areas originally included^13,136^. This step only affected the total number of parcels for the Craddock and Aicha parcellations of the human neocortex (i.e., fewer parcels than in the original volume representations).

### Brain phenotypes

We used various brain phenotypes specific to the human, macaque, marmoset, and mouse species to compare the parcel homogeneity of the geometric and existing benchmark parcellations. All data were obtained from published studies and/or open-access repositories, unless otherwise stated. All data were for the whole brain (includes left and right hemispheres), unless otherwise stated.

#### Human neocortex

We analyzed resting-state functional MRI (fMRI) FC matrices from three independent datasets [the Human Connectome Project (HCP)^14^; the Genomics Superstructure Project (GSP)^92^; and a dataset acquired in-house at Monash University (Monash)^93^] and 242 non-FC brain maps divided into the following 8 categories: morphometry (1 map); microstructure (3 maps); cytoarchitecture (50 maps); metabolism (5 maps); gene expression (1 map); chemoarchitecture (19 maps); HCP task activation (47 maps); and Neurosynth (116 maps). All data were mapped onto the fsLR_32k surface space, with 32,492 vertices per hemisphere.

The HCP FC matrix was derived from preprocessed resting-state fMRI data acquired in 255 unrelated healthy individuals (ages 22–35; 132 females), which is the largest HCP sample excluding twins or siblings and with all participants having completed resting-state and task-evoked data^14^. In brief, the fMRI parameters were: field strength of 3 T, isotropic voxel size of 2 mm, repetition time (TR) of 720 ms, echo time (TE) of 33.1 ms, left to right encoding direction, and scanning duration of 14.4 min with a total of 1200 time frames. All other acquisition parameters can be found in ref.^14^. Each individual’s data were preprocessed by the HCP team via their minimal preprocessing pipeline^142^, which included ICA-FIX to correct for structured noise and residual confounds^143^, and mapped onto the fsLR_32k surface space. For each participant, we calculated the Pearson correlation coefficient of pairs of vertex-level time series, resulting in a 32,492×32,492 FC matrix per hemisphere. We then took the average of the individual-specific FC matrices across the 255 individuals to construct a group-averaged FC matrix for each hemisphere.

The GSP FC matrix was derived from resting-state fMRI data acquired in 1564 healthy individuals (ages 18–35; 901 females)^92^. In brief, the fMRI parameters were: field strength of 3 T, isotropic voxel size of 3 mm, TR of 3000 ms, TE of 30 ms, and scanning duration of 6.2 min with a total of 124 time frames. All other acquisition parameters can be found in ref.^92^. Each individual’s data were preprocessed via the standard pipeline of fMRIPrep v22.0.0^144,145^, denoised with ICA-FIX, and mapped onto the fsLR_32k surface space. We then removed the first 5 time frames to stabilize the signal. We then calculated individual-specific FC matrices and took the average across the 1564 individuals to construct a group-averaged 32,492×32,492 FC matrix for each hemisphere.

The Monash FC matrix was derived from multiband resting-state fMRI data acquired in 440 healthy individuals (ages 21; all females)^93^. In brief, the fMRI parameters were: field strength of 3 T, isotropic voxel size of 3 mm, TR of 754 ms, TE of 21 ms, and scanning duration of 7.7 min with a total of 616 time frames. All other acquisition parameters can be found in ref.^93^. Each individual’s data were preprocessed via an in-house pipeline, which included ICA-FIX, global signal regression, and 3 mm spatial smoothing (see ref.^93^ for further details of the preprocessing), and mapped onto the fsLR_32k surface space. We then calculated individual-specific FC matrices and took the average across the 440 individuals to construct a group-averaged 32,492×32,492 FC matrix for each hemisphere. The diversity of preprocessing pipelines used across the HCP, GSP, and Monash datasets ensured that our findings are not specific to any single data processing approach.

The morphometry map represents cortical thickness and was obtained from the HCP repository^15^. The 3 microstructure maps represent the ratio of T1w and T2w MRI signal and diffusion MRI-derived estimates of neurite density index (NDI) and orientation dispersion index (ODI). T1w:T2w ratio indirectly quantifies intracortical myeloarchitecture and was obtained from ref.^98^, while NDI and ODI respectively quantify the packing density of axons or dendrites and orientational coherence of neurites and were obtained from ref.^146^. The 50 cytoarchitecture maps represent cell density profiles across 50 equidistant layers between the white and pial cortical layers of the BigBrain atlas^147^ and were obtained from ref.^148^.

The metabolism, gene expression, and chemoarchitecture maps were obtained from the neuromaps repository^96^. The 5 metabolism maps represent positron emission tomography (PET)-based markers of cerebral metabolic rate of glucose (CMRGlu), cerebral blood volume (CBV), cerebral blood flow (CBF), cerebral metabolic rate of oxygen (CMRO_2_), and synaptic density based on BPnd tracer binding to SV2a^149,150^. The gene expression map represents the first principal component of all genes in the Allen Human Brain atlas^151^. The 19 chemoarchitecture maps represent PET-based densities of neurotransmitter receptors of acetylcholine (α_4_β_2_, M_1_, and VACht), cannabinoid (CB_1_), dopamine (D_1_, D_2_, and DAT), GABA (GABA-A α_5_ subunit), glutamate (mGluR_5_ and NMDA), histamine (H_3_), mu-opioid (MOR), norepinephrine (NET), and serotonin (5-HT_1A_, 5-HT_1B_, 5-HT_2A_, 5-HT_4_, 5-HT_6_, and 5-HTT)^152–170^.

The 47 HCP task-activation maps represent unthresholded maps of task activations across 7 task domains of social, motor, gambling, working memory, language, emotion, and relational^171^. Each map represents a key contrast commonly used in the literature that shows the major activation pattern elicited by the task. See refs^47,171^ and Supplementary Table 3 for details of each task and contrast. We used the task maps as provided by HCP.

The 116 Neurosynth maps represent the association between voxels and cognitive terms obtained from Neurosynth, a meta-analytical tool that synthesizes results from more than 14,000 published fMRI studies by searching for keywords (e.g., action) published alongside fMRI voxel coordinates^97^. The unthresholded association maps were downloaded from https://neurosynth.org/. Although there are >1000 terms reported in Neurosynth, we only focused on 116 terms that are related to 11 cognition- and behavior-related concepts selected from the Cognitive Atlas^172^, which is a public ontology of concepts and terms used in cognitive science. See Supplementary Table 4 for the full list of terms we used.

#### Macaque neocortex

We analyzed 1 resting-state fMRI FC matrix and 19 non-FC brain maps divided into the following 3 categories: morphometry (1 map); microstructure (1 map); and chemoarchitecture (17 maps). All data were mapped onto the fsLR_10k surface space, with 10,242 vertices per hemisphere.

The FC matrix was derived from preprocessed resting-state fMRI data acquired in 6 individuals^173^. In brief, the fMRI parameters were: field strength of 3 T, isotropic voxel size of 1.25 mm, TRs ranging from 760 to 2600 ms, TEs ranging from 19 to 30 ms, and scanning duration of 40 min with a total of 1200 time frames. All other acquisition parameters can be found in ref.^173^. Each individual’s data were preprocessed via an HCP-style pipeline adapted to non-human primates (NHP) pipeline (https://github.com/Washington-University/NHPPipelines) and mapped onto the fsLR_10k surface space. We then calculated individual-specific FC matrices and took the average across the 6 individuals to construct a group-averaged 10,242×10,242 FC matrix for each hemisphere.

The morphometry and microstructure maps represent cortical thickness and T1w:T2w ratio, respectively, and were obtained from ref.^174^. The 17 chemoarchitecture maps represent autoradiographic densities in the left hemisphere of neurotransmitter receptors of acetylcholine M_1_, M_2_, and M_3_), adenosine 1, GABA (GABA-A, GABA-A/BZ, and GABA-B), glutamate (AMPA, Kainate, and NMDA), noradrenaline (α_1_ and α_2_), and serotonin (5-HT_1A_, 5-HT_2_) and averaged densities of excitatory (i.e., glutamate), inhibitory (i.e., GABA), and modulatory (i.e., serotonin, acetylcholine, and noradrenaline) receptors. The receptor maps were obtained from ongoing unpublished work, with all receptor density estimates reconstructed via the pipeline described in ref.^175^. Note that the chemoarchitecture maps were only available for the left hemisphere; hence, we restricted the calculation of homogeneity with respect to those maps to the left hemisphere.

#### Marmoset neocortex

We analyzed resting-state fMRI FC matrices from two independent sites [the Institute of Neuroscience (ION) in China and the National Institutes of Health (NIH) in the USA]^176^ and 3 non-FC brain maps divided into the following 3 categories: morphometry (1 map); microstructure (1 map); and cytoarchitecture (1 map). All data were mapped onto the fsaverage_38k surface space, with 37,974 and 38,094 vertices for the left and right hemispheres, respectively.

The FC matrices were derived from preprocessed resting-state fMRI data acquired in 10 individuals in each site (i.e., ION and NIH)^176^. In brief, the fMRI parameters for the ION site were: field strength of 9.4 T, isotropic voxel size of 0.5 mm, TR of 2000 ms, TE of 18 ms, right to left encoding direction, and scanning duration of 17.1 min with a total of 512 time frames. The fMRI parameters for the NIH site were: field strength of 7 T, isotropic voxel size of 0.5 mm, TR of 2000 ms, TE of 22.2 ms, right to left encoding direction, and scanning duration of 17.1 min with a total of 512 time frames. All other acquisition parameters can be found in ref.^176^. Each individual’s data were preprocessed via the Marmoset Brain Mapping Project’s pipeline (see https://marmosetbrainmapping.org/) and mapped onto the fsaverage_38k surface space. For each of the two sites, we then calculated individual-specific FC matrices and took the average across the 10 individuals to construct a group-averaged 37,974×37,974 and 38,094×38,094 FC matrix for the left and right hemispheres, respectively.

The morphometry and microstructure maps represent cortical thickness and T1w:T2w ratio, respectively, and were obtained from ref.^124^. The cytoarchitecture map represents cell density profiles in the left hemisphere derived from Nissl-stains and was obtained from ref.^177^. Note that the cytoarchitecture map was only available for the left hemisphere; hence, we restricted the calculation of homogeneity with respect to that map to the left hemisphere.

#### Mouse neocortex

We analyzed resting-state FC matrices derived from fMRI^178^ and Ca^2+^ imaging^179^ and 87 non-FC maps divided into the following 3 categories: microstructure (1 map); cytoarchitecture (1 map); and gene expression (85 maps). All data were mapped onto the Allen Mouse Brain CCFv3 volumetric space with 200 μm isotropic voxel resolution (except for the Ca^2+^ data, which were in 2D space; see below or Supplementary Information S4).

The fMRI FC matrix was derived from preprocessed resting-state fMRI data acquired in 193 individuals^178^. The data were pooled from 17 independent datasets across multiple centers. In brief, the fMRI parameters were: field strengths of 9.4 and 11.75 T, isotropic voxel size of 200 μm, TR of 1000 ms, TEs ranging from 10 to 25 ms, and number of time frames ranging from 150 to 1000. All other acquisition parameters can be found in ref.^178^. All data were preprocessed via a unified pipeline (see ref.^178^ for further details of the preprocessing) and mapped onto the CCFv3 volumetric space. We then calculated individual-specific FC matrices and took the average across the 193 individuals to construct a group-averaged voxel-level FC matrix for each hemisphere.

The Ca^2+^ FC matrix was derived from preprocessed resting-state cortex-wide fluorescence Ca^2+^ data acquired in 9 individuals^179^. In brief, the Ca^2+^ data were recorded using an in-house fiberscope^179^ and an sCMOS camera (512×512 pixels). The acquisition interleaved cyan (470/24, Ca^2+^-sensitive) and violet (395/25, Ca^2+^-insensitive) wavelengths using a Lumencor (LLE 7Ch Controller) light source at 20 Hz. The violet wavelength was for measuring background noise, which was regressed from the cyan data to obtain the final data at 10 Hz temporal resolution. The resulting images have a 25×25 μm resolution. The exposure time of each wavelength (violet and cyan) was 40 ms to avoid artifacts caused by the rolling shutter. All other acquisition parameters and preprocessing can be found in ref.^179^. Since the Ca^2+^ data were in 2D space, we projected the Allen Mouse Brain Atlas (i.e., ABA80 in Fig. 4A) and the corresponding geometric parcellation in 3D volume space to the 2D imaging plane of the Ca^2+^ data. See Supplementary Information S4 for further details about the imaging data, preprocessing, and volume-to-2D plane projection of the parcellations. We then calculated individual-specific FC matrices for each hemisphere. We used the pixel-level FC matrices and 2D plane-projected parcellations to calculate parcel homogeneity for each individual, which we then averaged for the final analysis.

The microstructure map represents T1w:T2w ratio and was obtained from refs^180,181^. The cytoarchitecture map represents cell density profiles derived from Nissl-stains and was obtained from ref.^182^. The 85 gene expression maps represent gene-expression profiles and were obtained from ref.^183^. Although there are >4000 gene-expression maps available from the above dataset, we restricted our analysis to 85 brain-related genes expressed in the brain chosen based on the criteria developed in ref.^181^. See Supplementary Table 5 for the list of genes we analyzed.

#### Human non-neocortical structures

We analyzed the same three independent resting-state fMRI datasets used for the human neocortex (i.e., HCP, GSP, and Monash). We analyzed the data in their original volumetric space smoothed with a full width at half maximum (FWHM) of 6 mm and then constructed a group-averaged voxel-level FC matrix for each hemisphere of each non-neocortical structure.

### Null FC matrices

In Figs 1C and 1E and Supplementary Figs 8A–B, we used the eigenstrapping method to randomize fMRI data and generate null FC matrices^91^ (see Supplementary Information S2.1). We first generated 1000 randomly rotated surrogate geometric eigenmodes of the human neocortical surface, separately for the left and right hemispheres (Supplementary Fig. 2A). We applied the surrogate eigenmodes to reconstruct each time volume of an individual’s fMRI data (Supplementary Fig. 2B), effectively creating 1000 surrogates of the vertex-by-time fMRI data. Note that the same set of rotated eigenmodes was used across individuals. For each surrogate, we calculated individual-specific FC matrices and took the average across the individuals to construct 1000 null group-averaged FC matrices for each hemisphere. This approach generated a randomized map that retains the exact spatial and temporal autocorrelation of the original data without relying on heuristics employed in other popular methods, such as the Spin test^95,184,185^, that could affect proper statistical inference on brain data^91,186,187^. We nonetheless replicated our null findings using the Spin test for completeness (see Supplementary Information S2.2 and Supplementary Fig. 8C).

### Random parcellations

In addition to the eigenstrapping and Spin test methods discussed in the previous section, we also calculated null FC homogeneity using 1000 random parcellations of the human neocortical surface (Fig. 1D). The parcellations were generated using in-house Matlab code to subdivide the cortical surface into *n* parcels with approximately equal surface area (Supplementary Figs 2C–D). The algorithm is as follows. An initial set of *n* seed vertices is first selected via farthest-point sampling based on geodesic distance to ensure wide coverage across the surface. Each vertex is then assigned to the nearest seed, producing an initial Voronoi-like pattern. We then performed multiple iterations to implement a Lloyd-like relaxation process^188^ to balance the surface area, where new centroids were computed for each parcel at each iteration. We implemented at most three iterations, because it provides a balance between computational accuracy and efficiency.

### Reaction-diffusion model

Reaction-diffusion models are used ubiquitously to study various forms of biological pattern formation^189^. The generalized model of two coupled molecules, *u* and *v*, spatiotemporally diffusing and reacting in a system is commonly defined in terms of the following equations^77,78^:

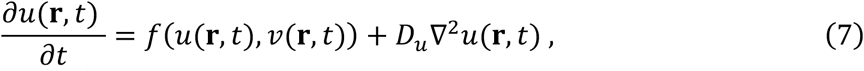

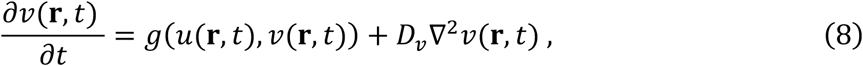

where *u*(**r**, *t*) and *v*(**r**, *t*) denote the local concentration of molecules *u* and *v*, respectively, at spatial location **r** at time *t*. For brevity, we will drop the (**r**, *t*) in subsequent equations, but *u* and *v* are treated as functions in space and time, unless otherwise stated. The terms *f*(*u*, *v*) and *g*(*u*, *v*) model the reactions of the molecules (e.g., production and elimination of concentrations), while the terms *D_u_*∇^2^*u* and *D_v_*∇^2^*v* model the diffusion of the molecules with *D_u_* and *D_v_* being positive diffusion constants. Assuming that the molecules have uniform concentrations at equilibrium, solutions that depart from the equilibrium can be obtained as follows (see Supplementary Information S1.1 for the derivation):

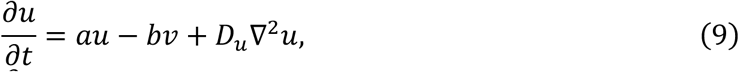

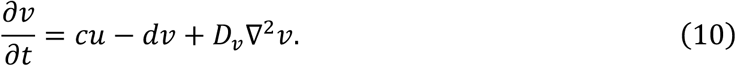

By setting the constants *a*, *b*, *c*, and *d* as positive numbers, the form of Eqs (9) and (10) assigns molecule *u* as the activator and molecule *v* as the inhibitor. Turing has shown that certain regimes of the model’s parameters (e.g., inhibitor diffuses much faster than the activator) can destabilize the uniform stationary states of the molecules’ concentrations, leading to the spontaneous emergence of a spatially non-constant solution at equilibrium that defines a spatial pattern, called a Turing instability^56^, that can mimic various patterns found in nature (e.g., leopard spots, zebra stripes; see refs^78,190^ for a collection of related articles). One can show that it is the relation between the model constants, and not their specific values, that gives rise to spatial Turing instabilities (see Supplementary Information S1.2 for the derivation). Specifically, instabilities will occur if the model constants satisfy all of the following conditions: 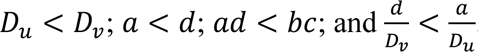.

In Fig. 7, we simulated the model in Eqs (9) and (10) on the left hemisphere of the human neocortex, seeding molecules *u* and *v* at distinct areas defined by the HCP-MMP1 atlas (i.e., Glasser360 in Fig. 2A)^2^. The example steady-state solution in Fig. 7B (top row) with Turing instability was obtained using the parameters: *a* = 2; *b* = 4; *c* = 2; *d* = 3; *D_u_* = 50; and *D_v_* = 100.

We reiterate that the spatial pattern of the instability shown in Fig. 7B is invariant to the specific values of these parameters as long as the model parameter combinations meet the conditions defined above for the instability to emerge. Moreover, our hierarchical framework need not necessarily result from a reaction-diffusion process, given prior work demonstrating that a bipartition could be obtained using a single diffusion gradient (as in Wolpert’s French Flag Problem^99,10^). This invariance to model parameters and dynamics supports the robustness of the underlying process of recursive bipartition based on geometry and suggests that multiple mechanisms may achieve the same end. We used an activator-inhibitor reaction-diffusion process here given evidence that some areal patterning molecules are mutually antagonistic (e.g., see refs^104,191,192^), but acknowledge that this is not the only process that may yield similar large-scale patterning. Indeed, experimental studies have shown that morphogen gradients can delineate regional tissue boundaries using a variety of physiological processes^66^ and that the actual processes driving the regional patterning of the brain are likely complex, involving time-dependent interactions between multiple molecular signals^52,63,193^. We welcome future work that improves the physiological plausibility of our model using additional processes such as those discussed above.

We further note that our analysis here focuses on determining how geometry constrains reaction-diffusion dynamics and early regional patterning, due to their formal relationship captured by the LBO in Eqs (9) and (10). Beyond patterning, geometry also plays an important role in brain function and connectivity. Under neural field theory^194^, geometry constrains wave-like dynamics^195,196^, which can explain many features of brain activity and function^47^. Furthermore, it can be shown that these geometric constraints prioritize a particular form of distance-dependent connectivity^186,197^ that dominates connectome organization^198,199^, and recent work has shown that it is possible to derive a generative model of connectome architecture purely from geometric eigenmodes^48^. Future work could explore how geometric constraints on areal patterning interact with activity-dependent and axonal pathfinding processes to shape the brain’s regional organization.

## Data availability

All source data to generate the results of the study and species-specific geometric parcellations of varying resolutions will be made openly available at https://github.com/NSBLab/BrainGeomParc upon publication.

## Code availability

All computer codes to generate the parcellations, analyze results, and reproduce the figures of the study will be made openly available at https://github.com/NSBLab/BrainGeomParc upon publication.

## Acknowledgements

This work was supported by Monash eResearch capabilities, including the M3 MASSIVE high-performance computing facility (www.massive.og.au). J.C.P. was supported by the Australian National Health and Medical Research Council (2034000), Monash Faculty of Medicine, Nursing, and Health Sciences (FMNHS) Early Career Postdoctoral Fellowship, and Monash FMNHS Early Career Research Excellence Program. M.M. was supported by the SNSF Postdoc Mobility Grant (214392). N.P.-G. was supported by the Helmholtz Association’s Initiative and Networking Fund through the Helmholtz International BigBrain Analytics and Learning Laboratory (HIBALL) under the Helmholtz International Lab grant agreement InterLabs-0015. B.T.T.Y. was supported by the NUS Yong Loo Lin School of Medicine (NUHSRO/2020/124/TMR/LOA), Singapore National Medical Research Council (NMRC) LCG (OFLCG19May-0035), NMRC CTG-IIT (CTGIIT23jan-0001), NMRC OF-IRG (OFIRG24jan-0030), NMRC STaR (STaR20nov-0003), Singapore Ministry of Health (MOH) Centre Grant (CG21APR1009), the Temasek Foundation (TF2223-IMH-01), and United States National Institutes of Health (R01MH133334). M.A.B. was supported by the National Health and Medical Research Council (1154378, 1146292, 1045354, 1006573). M.B. was supported by the National Health and Medical Research Council (2008612). A.F. was supported by the Australian Research Council (FL220100184), National Health and Medical Research Council (1197431), and Sylvia and Charles Viertel Charitable Foundation (2017042).

## Author contributions

J.C.P., P.A.R., K.M.A., and A.F. conceptualized the study. J.C.P., P.A.R., and A.F. designed the methodology. J.C.P. performed the investigation. M.M., X.S., and F.M. analyzed the mouse calcium imaging data. P.T.L., A.H., T.F., N.P.-G., J.T., M.A.B., R.T.C., and E.L. contributed data and data processing. P.A.R., R.K., B.T.T.Y., and M.B. provided intellectual input to data analysis. J.C.P. and A.F. administered the project. J.C.P. developed the visualizations. A.F. acquired funding. A.F. supervised the project. J.C.P. and A.F. wrote the original draft. All authors reviewed and edited the final manuscript.

## Competing interests

K.M.A. is the scientific advisor and a shareholder in BrainKey Inc., a medical image analysis software company. The other authors declare no competing interests.

## Notes

### Summary of Updates

Updated most sections of the main text; Figures 1 and 2 and Supplementary Figures 2, 3, and 8 revised

