## Supplementary Information for "Geometric influences on the regional organization of the mammalian brain"

### S1. Reaction-diffusion model

#### S1.1. Linearized equations

The generalized reaction-diffusion (RD) model of two coupled molecules, $u$ and $v$, is commonly defined in terms of the following equations[1,2]:

$$\frac{\partial u(\mathbf{r},t)}{\partial t} = f(u(\mathbf{r},t), v(\mathbf{r},t)) + D_u \nabla^2 u(\mathbf{r},t)\,, \qquad (S1)$$

$$\frac{\partial v(\mathbf{r},t)}{\partial t} = g(u(\mathbf{r},t), v(\mathbf{r},t)) + D_v \nabla^2 v(\mathbf{r},t)\,, \qquad (S2)$$

where $u(\mathbf{r},t)$ and $v(\mathbf{r},t)$ denote the local concentrations of molecules $u$ and $v$, respectively, at spatial location $\mathbf{r}$ at time $t$. For brevity, we will drop the $(\mathbf{r},t)$ in subsequent equations, but $u$ and $v$ are treated as functions in space and time, unless otherwise stated. The terms $f(u,v)$ and $g(u,v)$ model local dynamics (e.g., production and elimination of concentrations), while the terms $D_u\nabla^2 u$ and $D_v\nabla^2 v$ model diffusion dynamics with $D_u$ and $D_v$ being positive diffusion constants. We assume that an initial spatially homogeneous solution exists, $u = u_0$ and $v = v_0$, where $u_0$ and $v_0$ are constant equilibrium values. We can write $u$ and $v$ in terms of their departures, $\tilde{u}$ and $\tilde{v}$, from their equilibrium values as:

$$u = u_0 + \tilde{u}\,, \qquad (S3)$$

$$v = v_0 + \tilde{v}, \qquad (S4)$$

with $\tilde{u} \ll u_0$ and $\tilde{v} \ll v_0$. Therefore, Eqs (S1) and (S2) can be linearized around $\tilde{u}$ and $\tilde{v}$ to get:

$$\frac{\partial \tilde{u}}{\partial t} = a\tilde{u} - b\tilde{v} + D_u \nabla^2 \tilde{u}\,, \qquad (S5)$$

$$\frac{\partial \tilde{v}}{\partial t} = c\tilde{u} - d\tilde{v} + D_v \nabla^2 \tilde{v}\,, \qquad (S6)$$

where

$$a = \left.\frac{\partial f}{\partial u}\right|_{(u_0,v_0)}, b = -\left.\frac{\partial f}{\partial v}\right|_{(u_0,v_0)}, c = \left.\frac{\partial g}{\partial u}\right|_{(u_0,v_0)}, d = -\left.\frac{\partial g}{\partial v}\right|_{(u_0,v_0)}. \qquad (S7)$$

Assuming that the production constants $a$, $b$, $c$, and $d$ are positive, the signs in Eqs (S5) and (S6) designate $u$ as the activator and $v$ as the inhibitor. The linearized equations thus simulate the concentration dynamics of activator and inhibitor molecules, whereby the activator auto-catalytically enhances the production of both molecules while the inhibitor suppresses the growth. Note that Eqs (S5) and (S6) are the same as Eqs (7) and (8) of the Methods section of the main text, respectively, with the tilde above $u$ and $v$ removed for brevity.

#### S1.2. Turing instability

Turing's seminal work[3] showed that the spatially complex patterns generated by RD models are due to diffusion-driven instabilities, termed Turing instabilities. Here, we briefly show the standard derivation for the minimally sufficient conditions needed to induce Turing instability in Eqs (S5) and (S6), and we refer readers to ref.[1] for further details.

We first combine Eqs (S5) and (S6) in vector form as follows:

$$\frac{\partial \mathbf{W}}{\partial t} = \mathbf{AW} + \mathbf{D}\nabla^2\mathbf{W}\,, \tag{S8}$$

where

$$\mathbf{W} = \begin{pmatrix} \tilde{u} \\ \tilde{v} \end{pmatrix}, \qquad \mathbf{A} = \begin{pmatrix} a & -b \\ c & -d \end{pmatrix}, \qquad \mathbf{D} = \begin{pmatrix} D_u & 0 \\ 0 & D_v \end{pmatrix}. \tag{S9}$$

We make an ansatz that **W** has solutions of the form:

$$\mathbf{W} = \mathbf{W_0}\mathrm{e}^{\mathbf{ik\cdot r}+\sigma \mathrm{t}}\,, \tag{S10}$$

with wavevector **k**, growth rate σ, and complex constants $\mathbf{W_0}=\begin{pmatrix} \tilde{u}_0 \\ \tilde{v}_0 \end{pmatrix}$. Substituting Eq. (S10) into Eq. (S8) and rearranging terms, we get the eigenvalue problem:

$$(\mathbf{A} - \mathbf{D}\mathrm{k}^2)\mathbf{W_0} = \sigma\mathbf{W_0}\,, \tag{S11}$$

where $k = |\mathbf{k}|$. The characteristic polynomial to solve Eq. (S11) is:

$$\det[(\mathbf{A} - \mathbf{D}\mathrm{k}^2) - \sigma\mathbf{I}] = 0\,, \tag{S12}$$

$$\sigma^2 - \mathrm{tr}(\mathbf{A} - \mathbf{D}\mathrm{k}^2)\sigma + \det(\mathbf{A} - \mathbf{D}\mathrm{k}^2) = 0\,, \tag{S13}$$

where **I** is the 2×2 identity matrix, det denotes the determinant, and tr denotes the trace. The solutions of Eq. (S13) are:

$$\sigma = \frac{1}{2}\mathrm{tr}(\mathbf{A} - \mathbf{D}\mathrm{k}^2) \pm \frac{1}{2}\sqrt{[\mathrm{tr}(\mathbf{A} - \mathbf{D}\mathrm{k}^2)]^2 - 4\det(\mathbf{A} - \mathbf{D}\mathrm{k}^2)}. \tag{S14}$$

A Turing instability occurs when (i) there is a stable solution in the absence of diffusion and (ii) the solution becomes unstable when diffusion is added.

For condition (i), the absence of diffusion simplifies Eq. (S14) into:

$$\sigma = \frac{1}{2}\mathrm{tr}(\mathbf{A}) \pm \frac{1}{2}\sqrt{[\mathrm{tr}(\mathbf{A})]^2 - 4\det(\mathbf{A})}. \tag{S15}$$

Hence, a stable solution occurs when $\mathrm{tr}(\mathbf{A}) < 0$ and $\det(\mathbf{A}) > 0$, with $\mathrm{tr}(\mathbf{A}) = a - d$ and $\det(\mathbf{A}) = -ad + bc$. Therefore, the following relationships must be satisfied:

$$a < d \text{ and } \mathrm{ad} < \mathrm{bc}. \tag{S16}$$

For condition (ii), we analyze Eq. (S14) with diffusion included. Unstable solutions will occur only when $\det(\mathbf{A} - \mathbf{D}\mathrm{k}^2) > 0$, with:

$$\det(\mathbf{A} - \mathbf{D}\mathrm{k}^2) = (D_u k^2 - a)(D_v k^2 + d) + bc\,. \tag{S17}$$

The next step is to determine when Eq. (S17) becomes positive. Note that the shape of the function in Eq. (S17) is parabolic with respect to $k^2$ (parabola that opens upwards); hence, $\det(\mathbf{A} - \mathbf{D}\mathrm{k}^2) > 0$ will be achieved when the minimum at $k^2_{\mathrm{min}}$ is positive. We can find $k^2_{\mathrm{min}}$ by taking the derivative of Eq. (S17) with respect to $k^2$ to get:

$$k^2_{\mathrm{min}} = \frac{1}{2}\left(\frac{a}{D_u} - \frac{d}{D_v}\right). \tag{S18}$$

Equation (S18) will thus be positive when:

$$\frac{d}{D_v} < \frac{a}{D_u}. \tag{S19}$$

Finally, for the inequality in Eq. (S19) to be consistent with the first inequality in Eq. (S16), $D_u$ must be smaller than $D_v$ such that the inhibitor $v$ diffuses more rapidly than the activator $u$. Combining this relationship with those in Eqs (S16) and (S19), the four minimally sufficient conditions to induce a Turing instability are[1]: $D_u < D_v$; $a < d$; $ad < bc$; and $\frac{d}{D_v} < \frac{a}{D_u}$.

S1.3. Stability of the lowest-order non-constant geometric eigenmode

Mathematically, the spatial aspect of the RD equations in Eqs (S1) and (S2) is governed by the Laplacian operator, which appears in other fundamental physical equations such as those describing heat diffusion and wave dynamics[4]. The operator describes the curvature of the solution, which depends on the geometry of the medium within which the RD dynamics take place. At steady-state, the time-independent form of the RD equations in Eqs (S1) and (S2) reduces to the Helmholtz equation [i.e., Eq. (1) of the main text], which is precisely the equation from which the geometric eigenmodes are derived[5–7]. Hence, the emerging spatial pattern of any morphogen-like molecule diffusing continuously through the brain will naturally be constrained by the brain's geometry.

The wavelength of the Turing instability described in the previous section is set by the least stable and fastest-growing mode of the Laplacian, which is the lowest-order non-constant eigenmode, corresponding to the rostrocaudal geometric eigenmode with respect to the whole cortex. Instabilities with lower wavelengths are more stable and less likely to occur. Thus, the first pattern expressed in an RD process at the whole cortex level will coincide with the rostrocaudal axis, explaining why this axis is biologically preferred. This supports our use of the first non-constant geometric eigenmode, rather than other higher-order geometric eigenmodes with a single nodal line (i.e., patterns following the dorsoventral and mediolateral axes with respect to the whole cortex), as the simplest approximation of a biologically plausible geometry-constrained patterning process.

Our framework of partitioning a domain based on the lowest-order non-constant eigenmode also has a noteworthy analogy in graph theory, whereby the first non-constant eigenvector of a graph's Laplacian matrix (i.e., Fiedler vector) is the most efficient solution to partition a graph into two clusters while minimizing between-cluster weights and maximizing within-cluster weights[8]. Our framework is also analogous to spectral clustering techniques, which partition data arrays according to the dominant eigenvector[9–11]. We also note that while our formalism emphasizes geometric constraints on RD dynamics of morphogen-like substances, our framework can also capture the effects of activity-dependent processes in shaping regional organization to the extent that these processes are driven by diffusive dynamics. Under such scenarios, the dominant spatial modes of the dynamics also correspond to the dominant geometric eigenmodes, as dictated by the Helmholtz equation[6,7,12].

#### S1.4. Hierarchical dynamics and recursive bipartitions

In the main text and the sections above we have considered how our simple geometric framework is akin to an RD process at the coarsest possible scale—when the first division is made. We now consider how the recursive bipartitioning that occurs in our framework can be captured by a hierarchical RD process.

Following the first division of the brain into two subregions, concentrations of $u$ and $v$ will reach high levels at each of the subregions. We postulate that the high levels of $u$ and $v$ before saturation, and before higher-order eigenmodes destabilize, will trigger the RD process for two new sets of chemicals, $u_1$ and $v_1$ for subregion 1 and $u_2$ and $v_2$ for subregion 2. Without loss of generality, we assume that subregions 1 and 2 correspond to the subregion with high $u$ and high $v$, respectively, and the below discussion focuses on subregion 1. Analogously to Eqs (S1)–(S6), we can formulate the following related equations:

$$\frac{\partial \tilde{u}_1}{\partial t} = a_1\tilde{u}_1 - b_1\tilde{v}_1 + D_{u_1}\nabla^2\tilde{u}_1 , \qquad (S20)$$

$$\frac{\partial \tilde{v}_1}{\partial t} = c_1\tilde{u}_1 - d_1\tilde{v}_1 + D_{v_1}\nabla^2\tilde{v}_1 , \qquad (S21)$$

where $\tilde{u}_1$ and $\tilde{v}_1$ are departures of $u_1$ and $v_1$ from constant equilibrium values in the relevant subregion, respectively, $a_1$, $b_1$, $c_1$, and $d_1$ are the positive production constants, and $D_{u_1}$ and $D_{v_1}$ are the positive diffusion constants. We suppose that the production constants are proportional to $u$ such that at high $u$, the minimally sufficient conditions of a Turing instability will be achieved. Following our discussion in Section S1.3, a further Turing instability will evolve, leading to a new subdivision with spatial profile given by the lowest-order non-constant geometric eigenmode of the Laplacian within the subregion. Note that a similar instability occurs in subregion 2 where there is high $v$ and low $u$.

The above process can be repeated to achieve further hierarchical levels of subdivision. The occurrence of saturation would prevent more than 2-fold subdivision at a given level of hierarchy, while the onset of a further instability would produce the next level of subdivision. We reiterate that our goal is to show the theoretical plausibility of such a mechanism and motivate our geometrically constrained framework, not to identify the precise chemical or genetic molecules involved at each level. Nonetheless, one implication of our model is that the molecules are associated with the constants dictated by our model equations, ensuring that each successive level of subdivision is initiated before saturation is reached at the previous level. Termination of the hierarchy of subdivision could be achieved through three possible mechanisms: (i) concentration levels are lower than the threshold to induce destabilization of the modes of further subdivision; thresholds become higher as the subdivisions become smaller in size due to the general increase of the spatial wavenumber, $k$; (ii) activation of biological processes that release inhibitory signals for suppressing further differentiation, like the Hoxd13 genetic mechanism in digit formation[13], or the distinct, mutually inhibitory genetic programs that pattern sensory compared to association cortex[14]; and (iii) other processes that compete with the reaction-diffusion of patterning molecules, such as activity-dependent synaptic processes[15], becoming more dominant. Such termination could occur in a regionally specific manner, although we do not account for such variations in our framework for simplicity. Although one caveat of our current theoretical and simplified formulation is that it requires the identification of one unique molecule per subregion produced, $u$ and $v$ may represent sets of molecules, thereby allowing a combinatorically large number of possibilities. In fact, to produce $n$ subregions, we would only need about $\log_2 n$ hierarchical levels, which is biologically feasible.

### S2. Null models

#### S2.1. Eigenstrapping method for generating null data

In Figs 1C and 1E and Supplementary Figs 8A–B, we used the eigenstrapping method to randomize resting-state functional magnetic resonance imaging (fMRI) data and calculate null inter-vertex functional coupling (FC) homogeneity measures[16]. The eigenstrapping method leverages the mathematical properties of the geometric eigenmodes, derived from the Laplace-Beltrami operator of brain geometry [see Eq. (2) of the Methods section of the main text], to reconstruct any spatial map, $y(\mathbf{r})$, as a weighted sum of modes:

$$y(\mathbf{r}) = \sum_{j=1}^{M} \beta_j \psi_j(\mathbf{r}) + \epsilon(\mathbf{r}), \qquad (S22)$$

where $\beta_j$ is the amplitude of mode $j$ in explaining the data, which we calculated following ref.[6], $\psi_j$ is the $j$th mode, $M$ is the number of modes used, and $\epsilon$ is the residual error that vanishes when the full vector space of modes is used (e.g., total number of vertices $-$ 1 for a cortical surface mesh).

Geometric eigenmodes comprise spatial patterns with a specific spatial wavelength, which can be approximated using an idealized spherical case, following refs[6,7,17]. Due to the symmetry of a sphere, certain solutions are degenerate such that certain eigenmodes have the same eigenvalue and spatial wavelength, analogous to spherical harmonics in quantum physics. Hence, eigenmodes with similar spatial wavelength can be grouped into eigengroups (e.g., the first eigengroup comprises the rostrocaudal, dorsoventral, and mediolateral geometric eigenmodes with one nodal line). The eigenstrapping method uses this grouping by creating null eigenmodes, $\psi'$. In particular, for each eigengroup, the eigenmodes are projected from the brain's space to an abstract hypersphere, randomly rotated, and reprojected back onto the original space. This process is performed on all eigengroups to obtain $\psi'$ (Supplementary Fig. 2A). A surrogate version of the spatial map, $y'(\mathbf{r})$, is obtained by substituting the null eigenmodes in Eq. (S22) but using the original $\beta$ amplitudes and randomly permuting the residual error to preserve the power and variogram of the original data[16] (Supplementary Fig. 2B).

In our analysis, we generated 1000 surrogate geometric eigenmodes for each hemisphere, which we applied on each time volume of each individual participant's fMRI data, effectively creating 1000 surrogates of the vertex-by-time fMRI data that preserve both the spatial and temporal autocorrelation of the original data. Note that, for each surrogate instance, the same set of rotated modes was used across all time volumes and participants to ensure that later comparisons will be consistent. For each surrogate fMRI dataset, we calculated individual-specific FC matrices and took the average across the individuals to construct 1000 surrogate group-averaged FC matrices for each hemisphere. We applied the above process on our three resting-state fMRI datasets (i.e., HCP, GSP, and Monash), resulting in 1000 surrogate group-averaged FC matrices for each dataset. In Fig. 1C and Supplementary Fig. 8A, we compared the homogeneity of the geometric parcellations using the empirical and surrogate FCs. In Supplementary Fig. 8B, we compared the difference in homogeneity of the geometric and existing benchmark parcellations using the empirical and surrogate FCs. In Fig. 1E, we compared the local FC differentiation of the geometric and existing benchmark parcellations using the empirical and surrogate FCs.

S2.2. Spin method for generating null geometric parcellations

To further test whether the observed high FC homogeneity depends on the specific placement of the geometry-derived parcels, we used an alternative spatially constrained null model, called the Spin test[18–20], which rotates the geometric parcellations themselves rather than the FC data (Supplementary Fig. 8C). The Spin test projects the parcellation on a sphere, rotates the sphere by a random angle, and reprojects the parcellation back onto the surface space. Hence, the same rotation was applied to every vertex, such that the spatial features of the original parcellation were preserved but in a rotated frame. When the rotated vertices of the medial wall completely subsumed a parcel, that parcel was assigned a value of NaN and removed from subsequent homogeneity calculations, as per previous approaches[18–20]. This heuristic is not required for the eigenstrapping method. Also note that the Spin test can only be performed on surfaces, whereas the eigenstrapping method can be performed on both surfaces and volumes. We then generated 1000 surrogate geometric parcellations, which we used to calculate null FC homogeneity on the HCP, GSP, and Monash group-averaged resting-state FC matrices. In Supplementary Fig. 8C, we compared the FC homogeneity of the empirical and surrogate geometric parcellations.

### S3. Individual-specific geometric parcellations

Our main analyses in Figs 1–3 focused on creating parcellations for group-averaged templates of the human neocortex. However, we can directly apply our framework to individual cortical geometries[6,21,22], yielding parcellations that are specific to the individual's sulcal and gyral anatomy. In Supplementary Fig. 4, we generated individual-specific parcellations of varying resolutions using the midthickness surface of the 255 human participants from our HCP dataset (see 'Brain phenotypes' in Methods). Supplementary Figure 6A shows that using individual cortical geometries yielded parcel boundaries that better match the sulcal and gyral anatomy of the individuals compared to using the template surface. Note, however, that because of nuanced differences in individual geometries, it is difficult to establish corresponding parcels between individuals. Extending our framework to align parcels across participants is a topic for future investigation. However, Supplementary Fig. 4B shows that individual-specific geometric parcellations achieve more functionally homogeneous parcels than template geometric parcellations only at coarse resolutions (≤64 parcels). At finer resolutions, which is where most existing parcellations operate, individual-specific parcellations do not lead to significantly more functionally homogeneous parcels, suggesting that, for present purposes, the template-derived parcellations represent a good first approximation. Template-derived parcellations are also more suitable in our homogeneity calculations as many of our brain maps are from group-averaged data and are not specific to individuals.

To further show the flexibility of our framework in incorporating additional biological complexity, we implemented a hybrid version of our original approach. First, we noticed that at the specific scale of 16 parcels, the individual-specific geometric parcellations already capture aspects of visual and somatomotor fields (Supplementary Fig. 5A). We found that this effect is robust and highly reproducible across all individuals (Supplementary Fig. 5B). Moreover, these geometry-derived sensory fields overlapped with the counterpart parcels defined in the Glasser atlas and showed high functional homogeneity (Supplementary Figs 5B–D). Our hybrid approach manually 'locked' these putative sensory fields and only subdivided parts of the cortex that were outside these areas (i.e., no further subdivisions occurred within the putative sensory fields) until a desired number of parcels was reached, yielding a partition with stronger resemblance to classically defined areas (Supplementary Fig. 6A). However, the caveat to this hybrid approach is that it increases the variance of parcel sizes (Supplementary Fig. 6B), which reduces the global functional homogeneity of the atlas relative to the standard approach, especially at scales with ≥128 parcels (Supplementary Fig. 6C). This analysis nonetheless demonstrates how the framework can incorporate prior anatomical constraints when closer alignment with well-known cytoarchitectonic areas is desired.

### S4. Mouse calcium imaging data

#### S4.1. Data

We used mice data (n = 9) previously reported in refs[23,24]. Briefly, mice were 6–8 weeks old and 25–30 g at the time of the first imaging session. We included data from Slc17a7-cre/Camk2α-tTA/TITL-GCaMP6f (also known as Slc17a7-cre/Camk2α-tTA/Ai93) mice. Breeding and genotyping were done at Yale university, and all procedures were performed following the Yale Institutional Animal Care and Use Committee (IACUC). For permanent optical access to the cortical surface, all mice underwent a minimally invasive surgical procedure using an in-house built head-plate, as previously described[23,24].

S4.2. Imaging acquisition

Widefield calcium (WF-$Ca^{2+}$) imaging data were recorded using an in-house-built fiberscope[23]. The CamWare software version 3.17 and sCMOS cameras (512×512 pixels, pco.edge 4.2, PCO) were used. The images were relayed using 2,000,000 fiber optic cables (SCHOTT Inc.). Data acquisition entailed interleaving cyan (470/24, $Ca^{2+}$-sensitive) and violet (395/25, $Ca^{2+}$-insensitive) wavelengths using a Lumencor (LLE 7Ch Controller) light source at 20 Hz. The violet ($Ca^{2+}$-insensitive) wavelength was used as a measure of background noise and was regressed from the cyan data, resulting in a final (background noise corrected) 10 Hz temporal resolution. The images have a 25×25 μm resolution. The exposure time of each wavelength (violet and cyan) was 40 ms to avoid artifacts caused by the rolling shutter. Thus, the sequence was: 10 ms blank, 40 ms violet, 10 ms blank, 40 ms cyan, and so on.

S4.3. Preprocessing

Imaging data were registered to an in-house template space, previously described in ref.[25]. This template was then registered to the Allen Atlas reference space CCFv3[26]. Here, we used the in-house template space to generate the 40-parcel geometric parcellation per hemisphere.

The WF-$Ca^{2+}$ imaging data wavelengths ($Ca^{2+}$-sensitive and $Ca^{2+}$-insensitive) were separated and processed in parallel. Data were smoothed (sigma 0.1 mm), and 50 frames were discarded from the beginning and end of each run. The data were downsampled and noise ($Ca^{2+}$-insensitive) was regressed from the signal ($Ca^{2+}$-sensitive). $\Delta F/F_0$ was computed, and the data were filtered at 0.008–0.2 Hz. The $\Delta F/F_0$ timeseries was registered to the common space template by concatenating two linear transformations: (1) WF-$Ca^{2+}$ images → Angiogram; and (2) Angiogram → Multi-spin-multi-echo (MSME). These were both generated by hand using tools designed for this purpose within the BioImage Suite (BIS), as described previously[23].

At the conclusion of these steps, the data from all mice and imaging sessions reside in the in-house template space where all subsequent analyses were performed.

S4.4. Projecting the parcellations onto a 2D plane

In our homogeneity analyses, we compared the Allen Atlas (ABA80) and corresponding geometric parcellation (geometric80), both having 40 parcels per hemisphere and in three-dimensional (3D) MRI voxel space. In both cases, we limited the analysis to parcels visible in the field of view (FOV) of the cortical surface (i.e., excluding parcels below the cortex or along the sides of the brain, which are both not fully visible from above). For ABA80, only 30 parcels per hemisphere were fully visible, while for geometric80, 33 parcels per hemisphere were fully visible. Hence, we calculated parcel homogeneity only on visible parcels in each atlas. As the WF-$Ca^{2+}$ imaging data were 2D, we created a 'surface projection' of the 3D atlases using custom tools within BIS[23]. Briefly, the parcel labels were projected along the translation perpendicular to the optical imaging plane by shooting 'rays' from 'above'. Each ray was followed until it 'hits' the MRI brain-volume 'surface'. For each voxel in the MRI image, we took the parcel label at the surface of the MRI volume. This procedure enabled moving labels generated in 3D MRI space to the 2D imaging plane of the WF-$Ca^{2+}$ imaging data.

### S5. Supplementary Figures

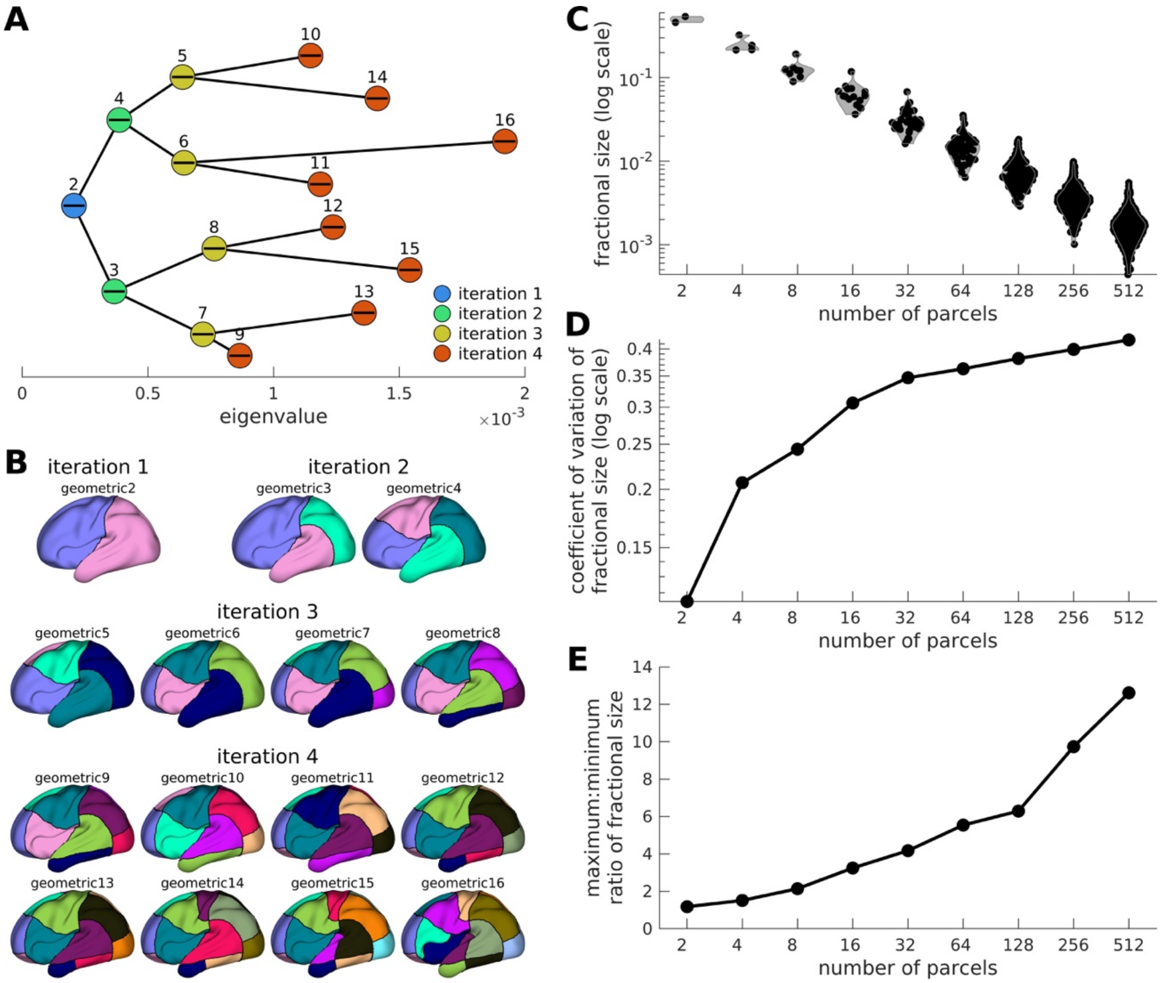

**Supplementary Fig. 1. Hierarchical bipartitioning method applied to the left human neocortex.** (**A**) Eigenvalue tree used to order the subdivisions. To construct the tree, we performed our hierarchical bipartitioning method up to 9 iterations (only the first 4 iterations are shown for brevity), storing the first non-constant geometric eigenmode and its corresponding eigenvalue for every division. We then arranged the divisions according to the magnitude of the eigenvalue (circles) but keeping track of which iteration the division belongs to (colors) and the parent-daughter subdivisions (lines). Depending on the desired parcellation resolution, the order of subdivisions in the eigenvalue tree is followed, prioritizing the subdivision with the lowest eigenvalue. Note that all subdivisions within an iteration must be chosen first before moving to the next iteration. The numbers above the circles show the number of parcels generated upon choosing that subdivision and represent how one should follow the trail along the tree. (**B**) Multiscale geometric parcellations with 2 to 16 parcels. The number beside the parcellation's name denotes the number of parcels for one hemisphere. (**C**) Distribution of parcel sizes for geometric parcellations with $2^N$ ($N$ = 1 to 9) number of parcels. The y-axis shows the fractional size of each parcel (i.e., number of vertices in each parcel relative to the total number of vertices in the hemisphere). (**D**) Coefficient of variation of parcel sizes. (**E**) Ratio of the maximum and minimum parcel sizes, approximating the compactness of the parcellation. Results in panels C to E show that higher resolutions of the geometric parcellations lead to smaller parcels but with more varied sizes.

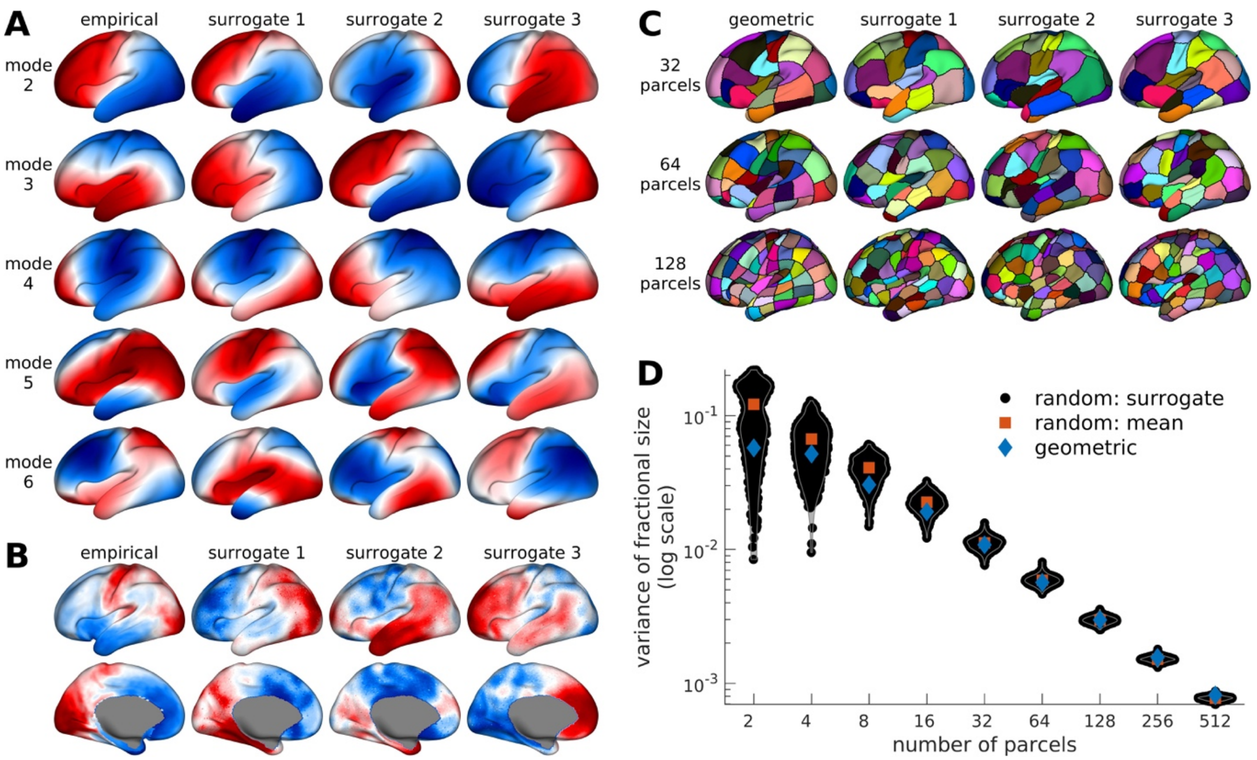

**Supplementary Fig. 2. Eigenstrapping and random parcellation null models applied to the human neocortex.** (**A**) Empirical and example surrogate eigenmodes of the left hemisphere generated by the eigenstrapping method. For brevity, only modes 2 to 6 and three example surrogates are shown. (**B**) Empirical and example surrogate maps of the left hemisphere. (**C**) Geometric and example surrogate random parcellations of the left hemisphere with 32, 64, and 128 parcels (from top to bottom). The random parcellations are obtained by randomly dividing the cortex into parcels of approximately equal surface area. (**D**) Distribution of parcel sizes for geometric parcellations (blue diamonds) and 1000 random parcellations (black dots) of the left hemisphere with $2^N$ ($N$ = 1 to 9) number of parcels. The y-axis shows the fractional size of each parcel (i.e., number of vertices in each parcel relative to the total number of vertices in the hemisphere). The red squares represent the means of the surrogate distributions. The panel shows that the geometric and random parcellations have broadly similar spread of parcel sizes.

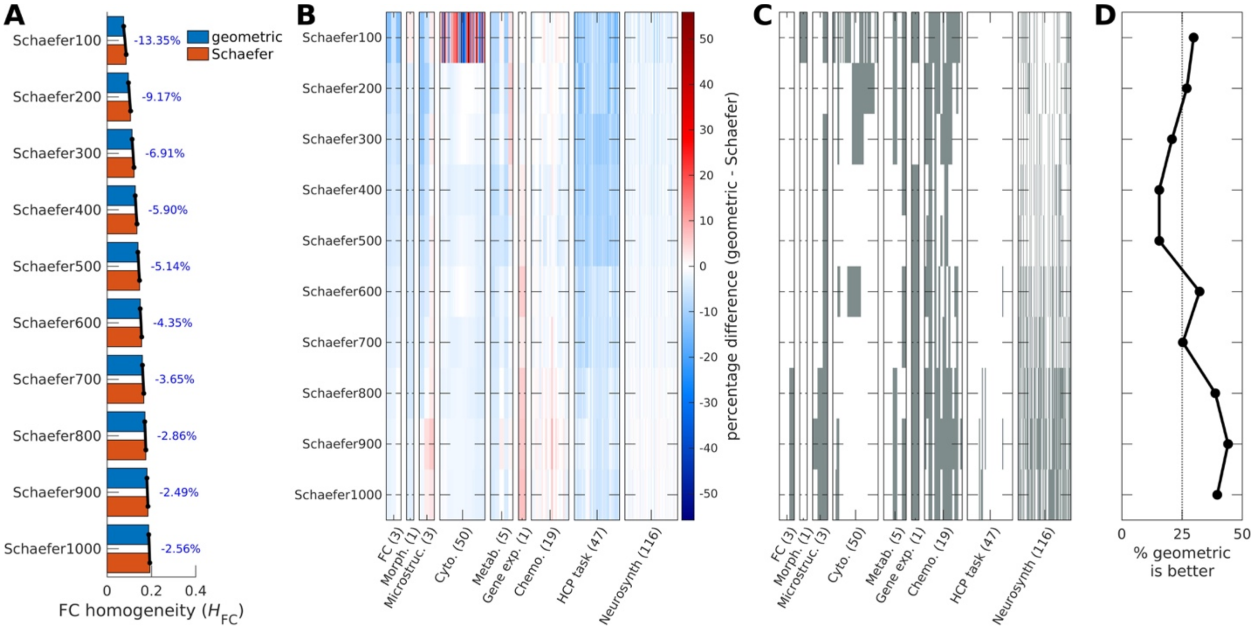

**Supplementary Fig. 3. Homogeneity of geometry-derived parcels of the human neocortex relative to different scales of the Schaefer parcellation.** (**A**) FC homogeneity ($H_{FC}$) calculated from the HCP dataset. The number beside the parcellation's name denotes the total number of parcels across both hemispheres (e.g., Schaefer100 has 100 parcels in total across the left and right hemispheres). The numbers beside the bar graphs represent the percentage difference between the geometric and Schaefer parcellations, where negative/blue indicates higher homogeneity for the Schaefer parcellations. (**B**) Percentage difference in homogeneity between geometric and Schaefer parcellations. Each row shows the result for one benchmark comparison and each column shows one brain map (including the results in panel A), where red indicates higher homogeneity for geometric parcellations. (**C**) Binarized version of panel B, where gray indicates higher or equivalent homogeneity for geometric parcellations, with equivalence quantified as a difference threshold of ±1%. (**D**) Percentage of maps for which geometric parcellations show higher or equivalent homogeneity. These results show that differences in homogeneity between the geometric and Schaefer parcellations attenuate from coarse to fine spatial scales.

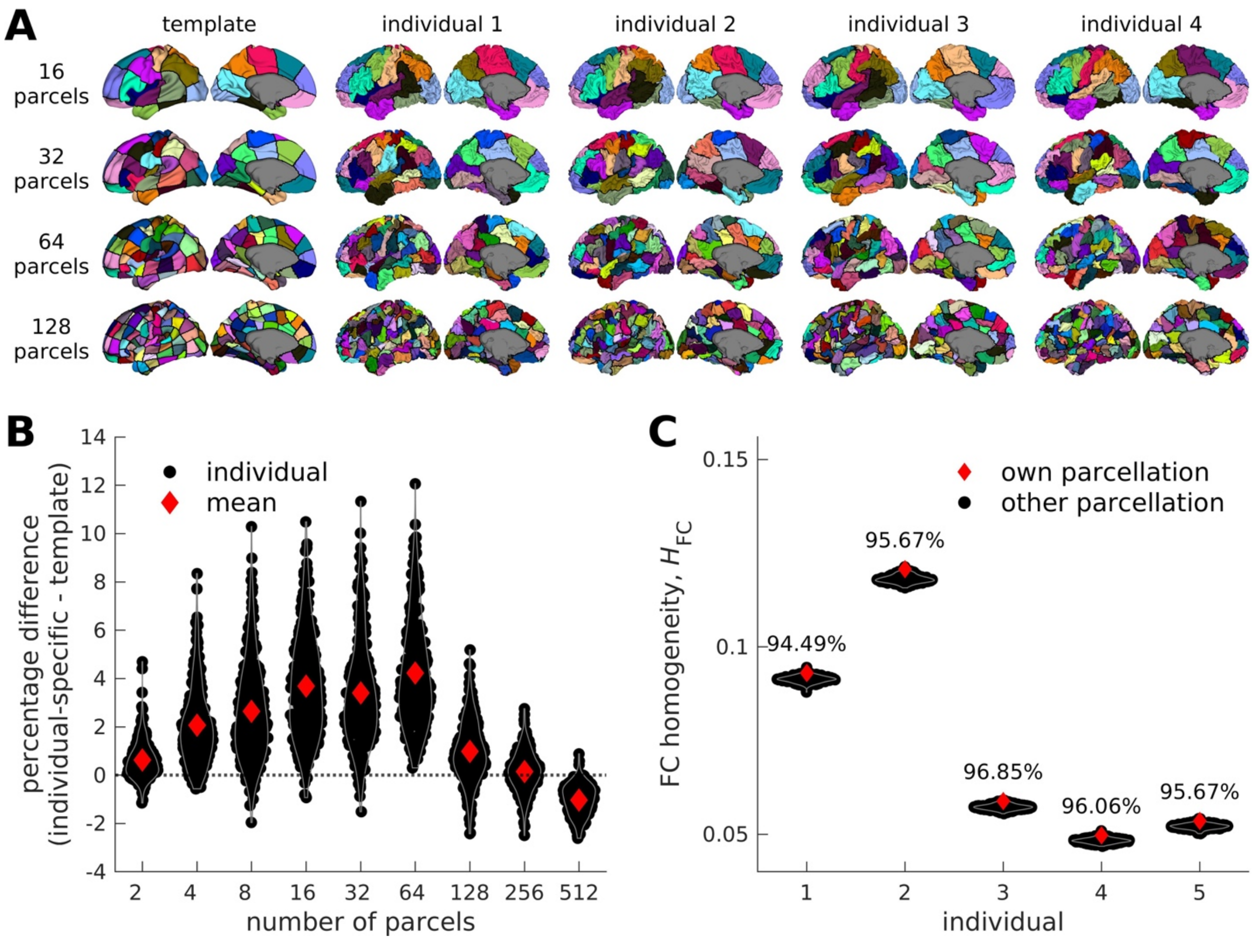

**Supplementary Fig. 4. Individual-specific geometric parcellations of the left human neocortex.** (**A**) Template-derived and example individual-specific geometric parcellations of four participants from the HCP dataset with 16, 32, 64, and 128 parcels (from top to bottom). The parcellations are shown on the template and the individual's midthickness surface to better demonstrate that the boundaries of the individual-specific geometric parcellations align with macroscale anatomical features (e.g., sulci). (**B**) Distribution of percentage differences in FC homogeneity ($H_{\mathrm{FC}}$) of individual-specific geometric parcellations relative to template geometric parcellations for different number of parcels. Each dot represents an individual and the red diamonds represent the means of the distributions. Individual-specific parcellations have generally higher homogeneity, especially for coarse resolutions. (**C**) Specificity of individual-specific geometric parcellations with 64 parcels for five example individuals from the HCP dataset. For each individual's FC data, we calculated $H_{\mathrm{FC}}$ using their own individual-specific parcellation (red diamonds) and using parcellations specific to 254 other individuals (black dots). The percentage above the distributions shows how many times an individual's own parcellation have higher $H_{\mathrm{FC}}$ than other individuals' parcellation. The high values demonstrate the specificity of the individual-specific parcellations.

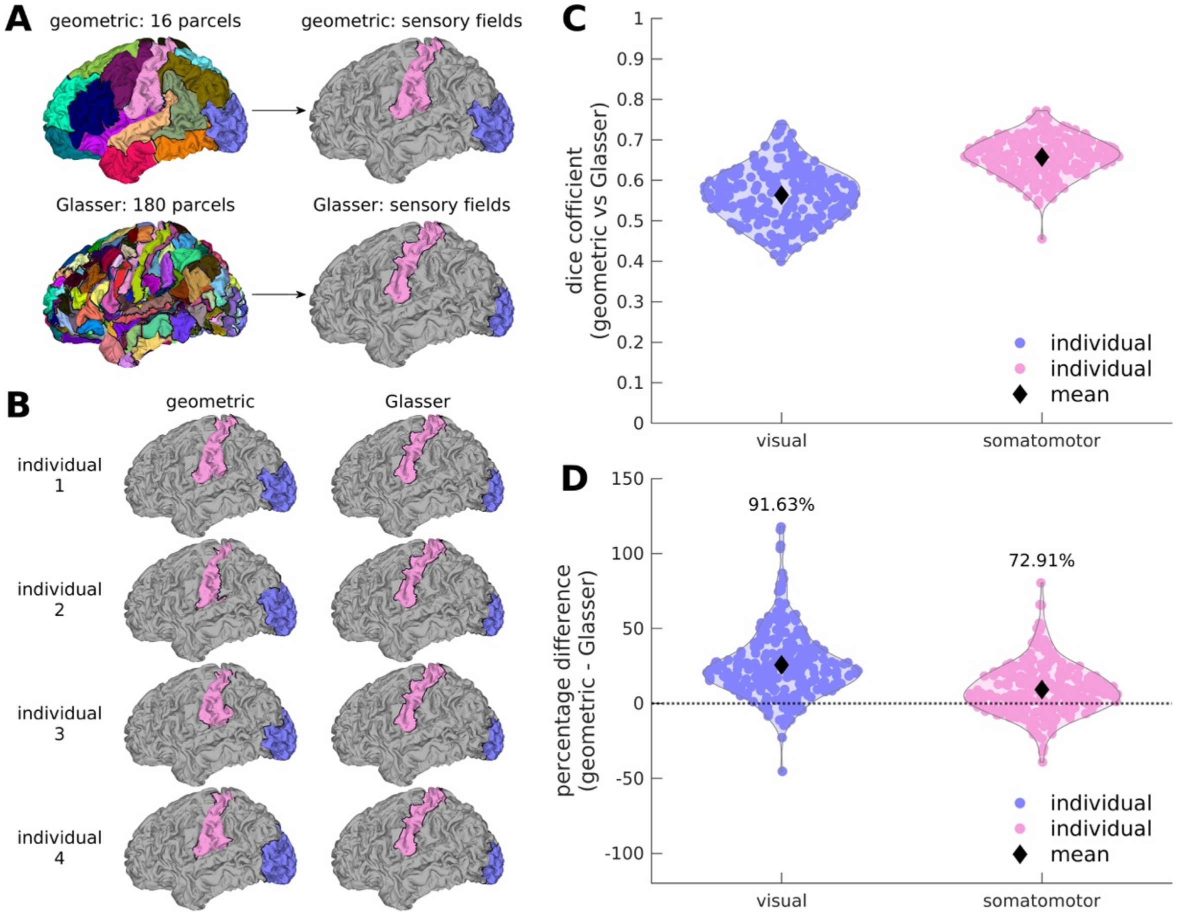

**Supplementary Fig. 5. Individual-specific geometric parcellations of sensory fields of the left human neocortex.** (**A**) Parcels resembling visual (blue) and somatomotor (pink) fields are directly extracted from the (Top) individual-specific geometric parcellations in Supplementary Fig. 4A at the scale of 16 parcels and (Bottom) Glasser atlas. (**B**) Geometric (left column) and Glasser (right column) parcellations of sensory fields in four example individuals of the HCP dataset. (**C**) Distribution of Dice coefficients between the visual and somatomotor fields of the geometric and Glasser parcellations across 255 individuals. Each dot represents an individual and the black diamonds represent the means of the distributions. There is a high degree of overlap between the geometric and Glasser parcellations. (**D**) Distribution of percentage differences in FC homogeneity ($H_{\mathrm{FC}}$) of individual-specific geometric parcellations relative to Glasser parcellations of visual and somatomotor fields. Each dot represents an individual and the black diamonds represent the means of the distributions. The percentage above the distributions shows how many times the geometric parcellation has higher homogeneity than the Glasser parcellation. The high values demonstrate the success of our framework in simultaneously capturing functionally homogeneous parcels that mimic anatomically known cortical areas.

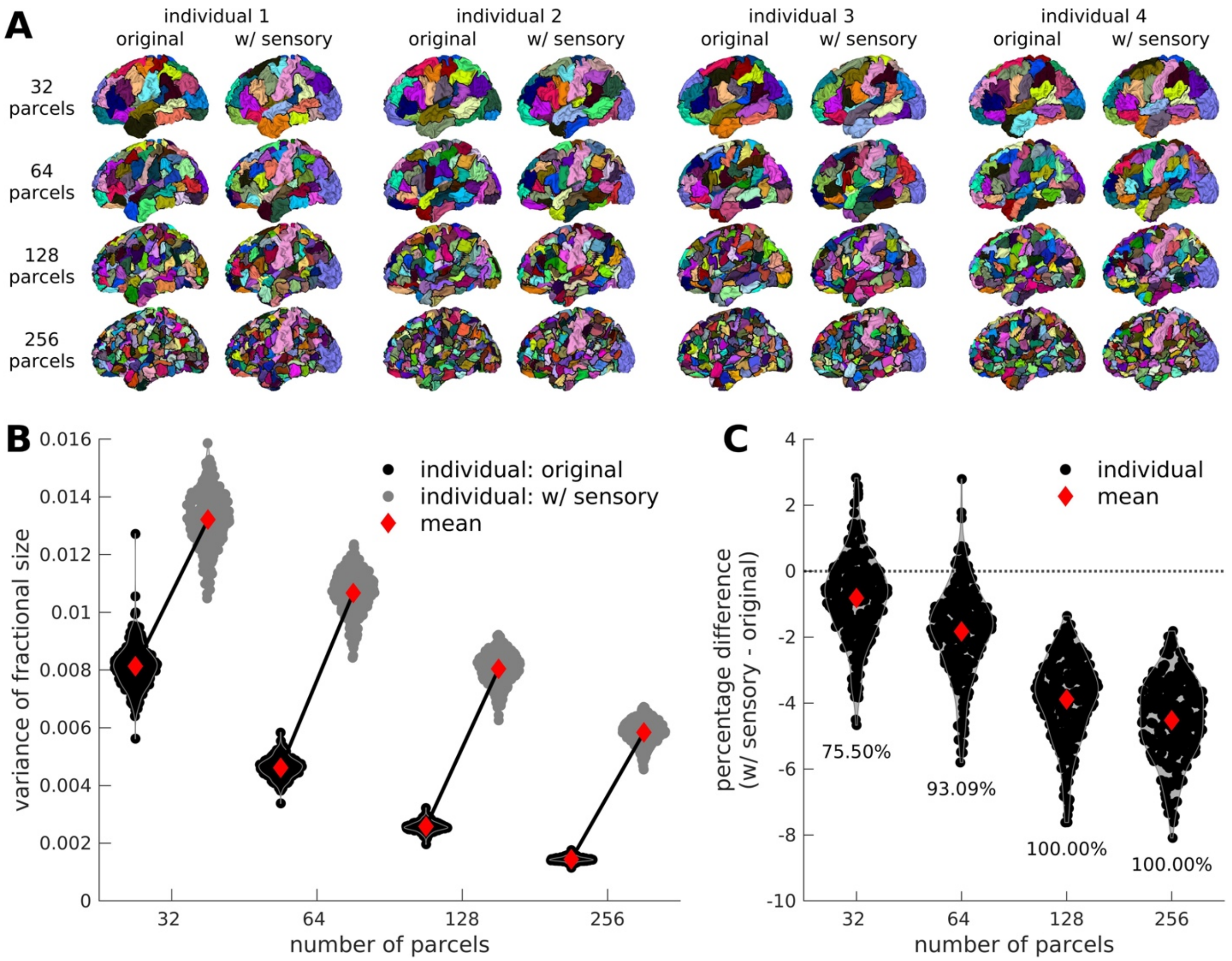

**Supplementary Fig. 6. Original and hybrid individual-specific geometric parcellations of the left human neocortex.** (**A**) Individual-specific geometric parcellations generated using our original approach ('original') in Supplementary Fig. 4A and a hybrid approach ('w/ sensory') where we first manually 'lock' the sensory fields in Supplementary Fig. 5B before further subdividing the rest of the cortex. Each column shows parcellations for an individual with 32, 64, 128, and 256 parcels (from top to bottom). (**B**) Distribution of parcel sizes for geometric parcellations generated using the original (black dots) and hybrid (gray dots) approaches with 32, 64, 128, and 256 parcels. The red diamonds represent the means of the distributions. (**C**) Distribution of percentage differences in FC homogeneity ($H_{\mathrm{FC}}$) of individual-specific geometric parcellations generated using the hybrid approach relative to the original approach with 32, 64, 128, and 256 parcels. Each dot represents an individual and the red diamonds represent the means of the distributions. The percentage below the distributions shows how many times the original approach has higher homogeneity than the hybrid approach. The results demonstrate that manually locking parcels, such as the sensory fields, impedes the homogeneity of the geometry-derived parcels.

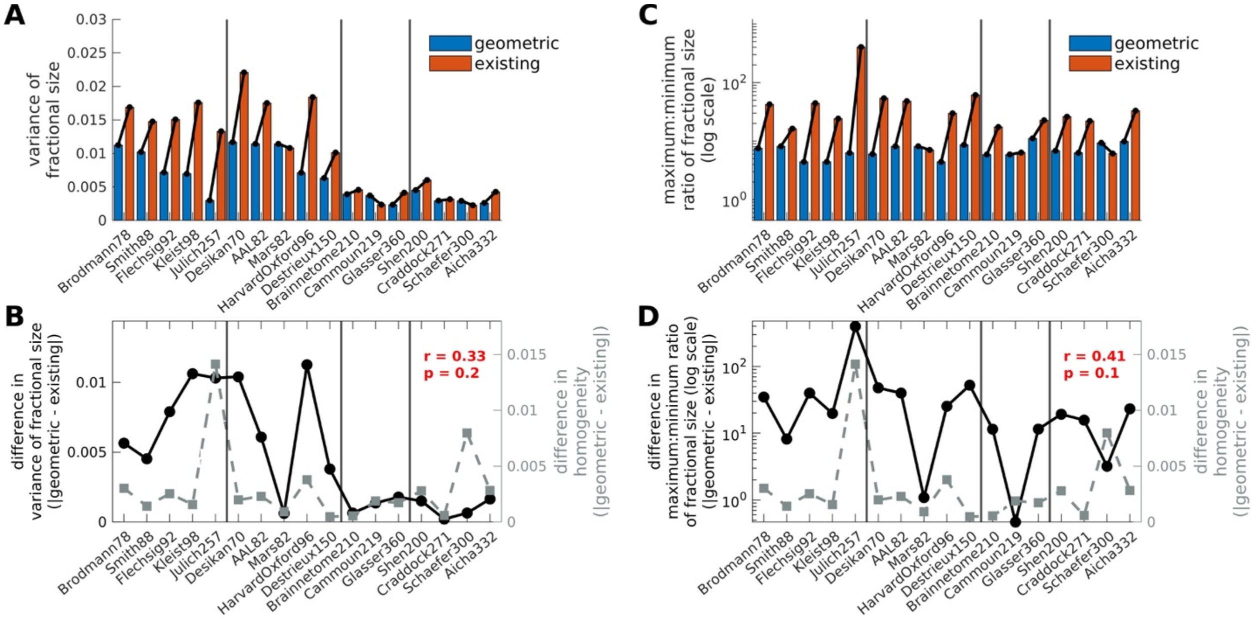

**Supplementary Fig. 7. Comparison of homogeneity and parcel sizes of geometric and existing benchmark parcellations of the left human neocortex.** (**A**) Variance of fractional parcel sizes, showing that geometric parcellations have parcel sizes that are generally more uniform than those of their counterpart existing parcellations. The only exceptions are Mars82, Cammoun219, and Schaefer300. (**B**) Absolute difference in the variance of fractional parcel sizes (left y-axis, circles, and solid line) and FC homogeneity (right y-axis, squares, and dashed line) between geometric and benchmark parcellations. The homogeneity was calculated using the HCP resting-state fMRI dataset. The effect size is small and not statistically significant ($r$ = 0.33, $p$ = 0.2), indicating that the association between differences in homogeneity and parcel size variance between the geometric and existing parcellations are not better by chance. (**C**) Same as panel A but for the ratio of the maximum and minimum parcel sizes, showing that geometric parcellations are more compact relative to their counterpart existing parcellations. (**D**) Same as panel B but the left y-axis shows the absolute difference in the ratio of the maximum and minimum fractional parcel sizes. The effect size is small and not statistically significant ($r$ = 0.41, $p$ = 0.1), indicating that the association between differences in homogeneity and parcel size compactness between the geometric and existing parcellations are not better by chance.

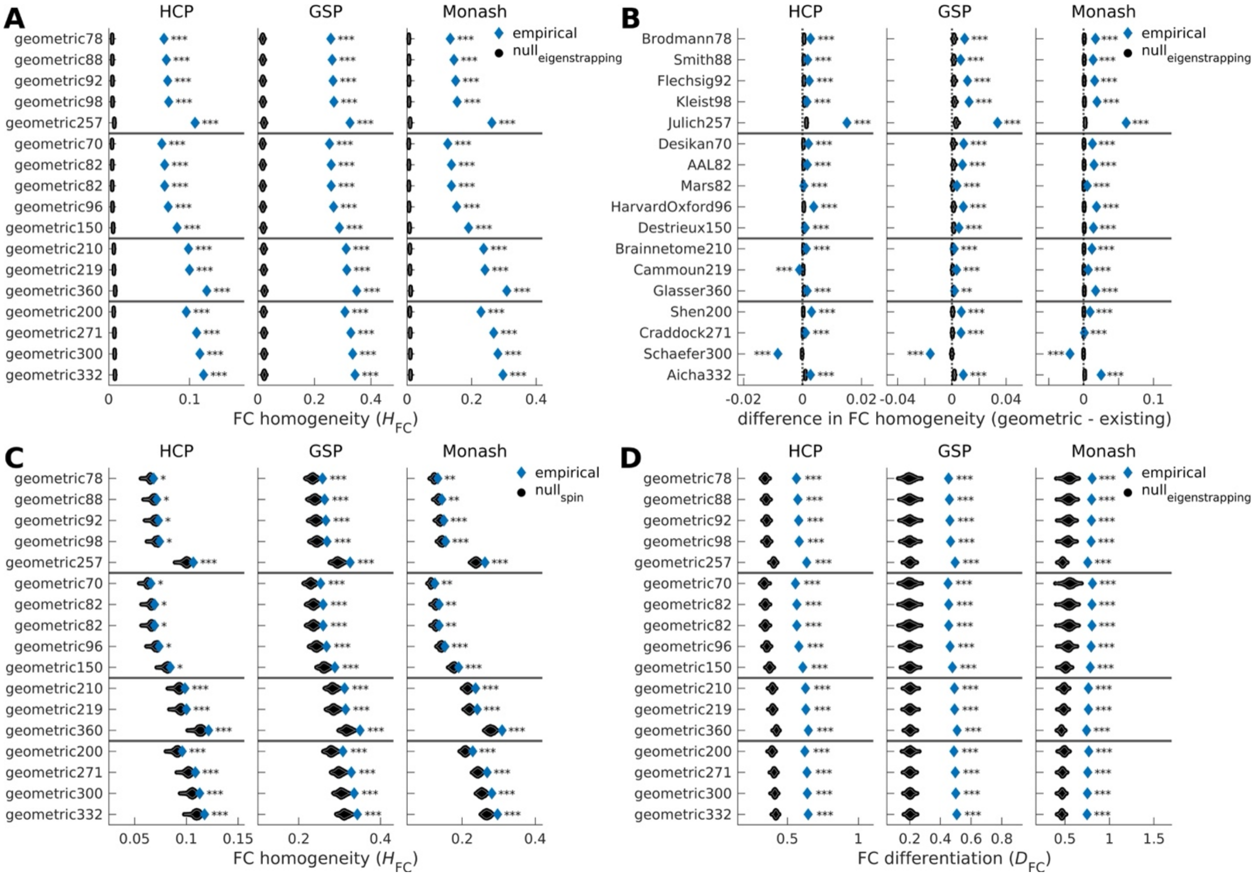

**Supplementary Fig. 8. Comparison of empirical and null FC homogeneity and differentiation of parcellations of the human neocortex across both hemispheres.** (**A**) Empirical and null FC homogeneity ($H_{\mathrm{FC}}$) of the counterpart geometric parcellations in Fig. 2A based on 1000 randomized FCs generated by the eigenstrapping method. (**B**) Empirical and null difference in FC homogeneity between geometric and benchmark parcellations based on 1000 eigenstrapping nulls. (**C**) Same as panel A but the nulls were based on 1000 spun versions of the parcellations. (**D**) Same as panel A but for local FC differentiation ($D_{\mathrm{FC}}$). For all panels, the blue diamonds correspond to the empirical data and the black dots correspond to the null data. Each geometric parcellation showed significantly higher empirical $H_{\mathrm{FC}}$ and $D_{\mathrm{FC}}$ than the null data with varying levels of significance (***: two-sided p-value <0.001; **: two-sided p-value <0.01; *: two-sided p-value <0.05).

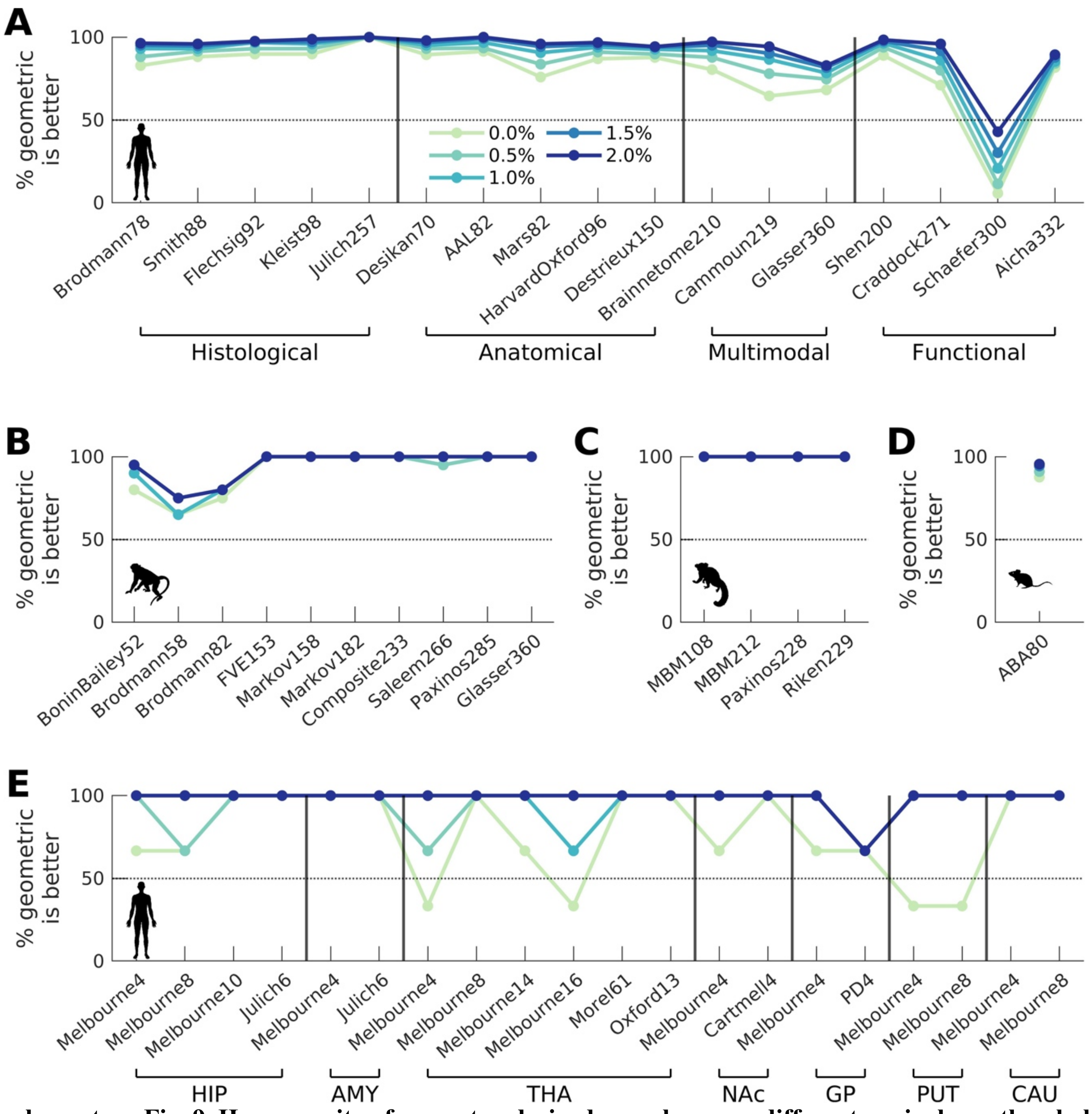

**Supplementary Fig. 9. Homogeneity of geometry-derived parcels across different equivalence thresholds.** The panels show the percentage of maps where the geometric parcellations have higher or equivalent homogeneity than the existing parcellations for the (**A**) human neocortex, (**B**) macaque neocortex, (**C**) marmoset neocortex, (**D**) mouse neocortex, and (**E**) human non-neocortical structures. The percentage was calculated using different definitions of equivalence in the homogeneity difference. The equivalence thresholds are 0.0%, 0.5%, 1.0%, 1.5%, and 2.0% (from light to dark colors).

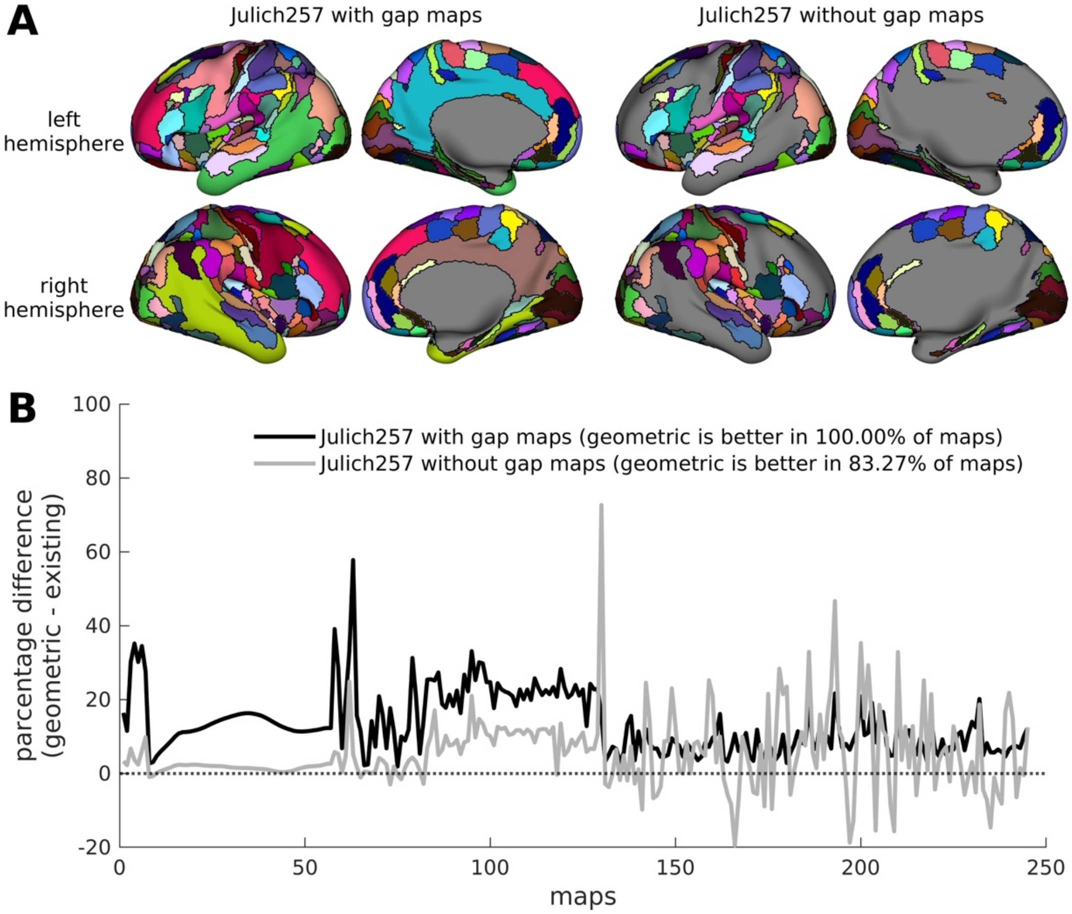

**Supplementary Fig. 10. Homogeneity of the Julich257 atlas with and without the gap maps.** (**A**) Julich257 atlas across both hemispheres with (left column) and without (right column) the gap maps. The gap maps represent areas deemed by Julich's approach to have ambiguous identities[27]. (**B**) Percentage difference in homogeneity between geometric parcellations and Julich257 atlas with (black line) and without (gray line) the gap maps across all FC and non-FC phenotypes or maps. Note that we removed all vertices within the gap maps in the geometric parcellations before calculating the homogeneity for fair comparison. The results demonstrate that geometric parcellation remains to have higher homogeneity (i.e., better than >83.27% of the maps) regardless of which version of the Julich atlas is used.

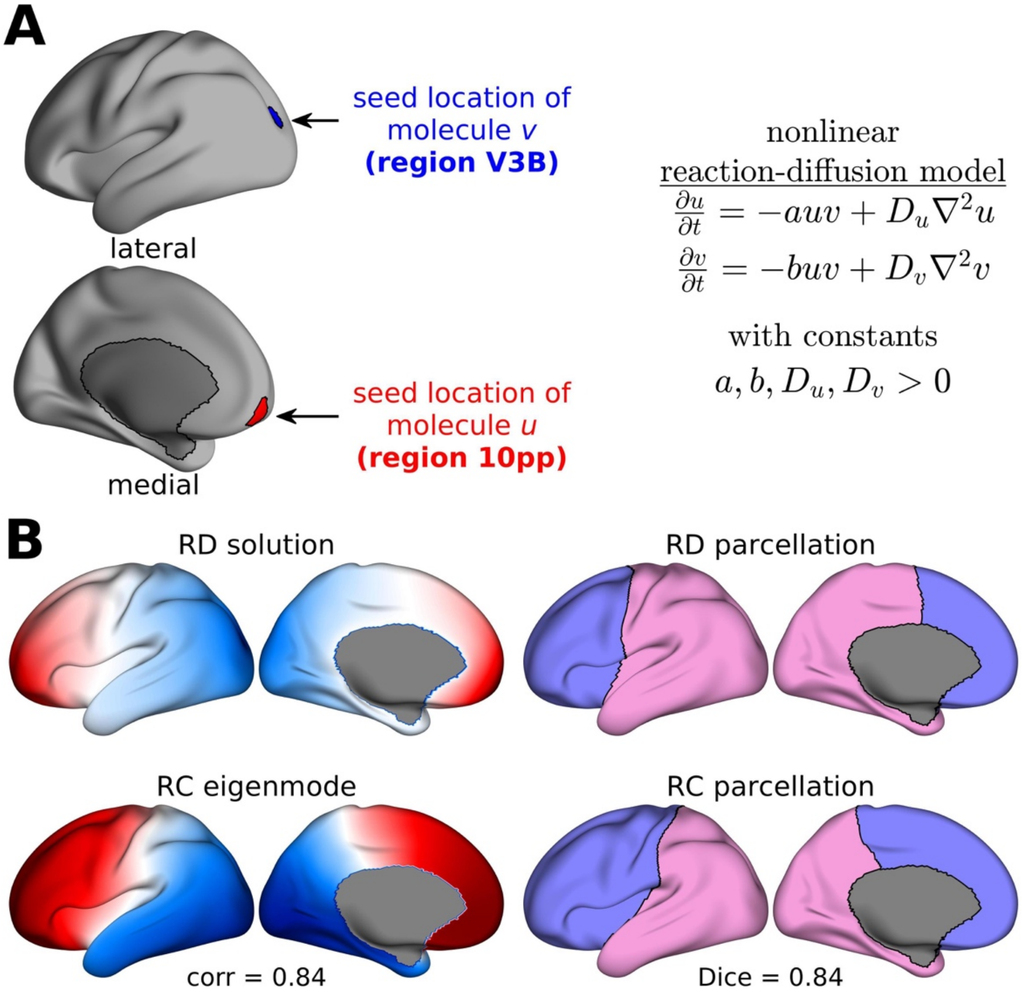

**Supplementary Fig. 11. Nonlinear reaction-diffusion model of regional organization.** (**A**) We used a nonlinear RD model (compared to the linear model in Fig. 7A) that assumes two molecules, $u$ and $v$, are seeded from distinct locations and diffuse in space and time with diffusion constants $D_u$ and $D_v$, respectively. Following Fig. 7A, we seeded the molecules at maximally distant points near the rostral and caudal poles, specifically at regions 10pp (orbital prefrontal cortex) and V3B for $u$ and $v$, respectively, based on the Glasser360 parcellation[28]. The constants $a$, $b$, $D_u$, and $D_v$ are assumed to be positive. All parameter values are set to the values of the same variables in Fig. 7. (**B**) The top row shows an example model RD solution ($u - v$) and the corresponding binary subdivision when the molecules are seeded at the locations described in panel A. The bottom row shows the rostrocaudal (RC) geometric eigenmode and the corresponding binary subdivision. Correspondence between the model RD solution and RC eigenmode is quantified using spatial correlation, and correspondence between the binary subdivisions is quantified using Dice coefficient. These results demonstrate the robustness of our findings to the choice of RD model type.

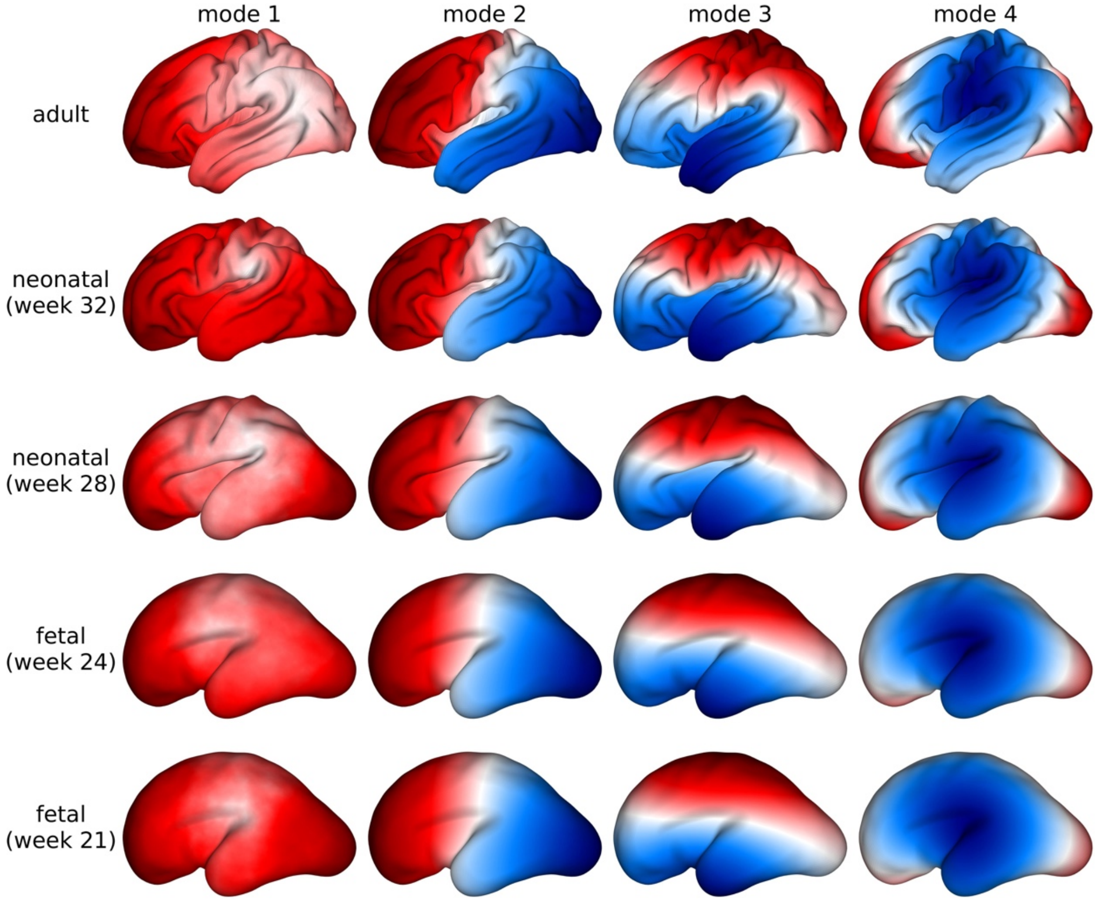

**Supplementary Fig. 12. Low-order geometric eigenmodes of human adult, neonatal, and fetal cortices.** Columns from left to right show the first four low-order geometric eigenmodes. Rows from top to bottom show the eigenmodes for a young adult brain, a 32-week neonatal brain, a 28-week neonatal brain, a 24-week fetal brain, and a 21-week fetal brain, respectively. The neonatal brains were obtained from postnatal imaging, while the fetal brains were obtained from in utero imaging. Both the neonatal and fetal cortical surface reconstructions were downloaded from the Developing Human Connectome Project[29]. The results show that the low-order modes across ages show high spatial correspondence. Note that mode 2 is used as the starting point for our framework and is largely invariant, suggesting that our reliance on adult models offers a reasonable first approximation of geometric constraints on the developmental process influencing regional organization.

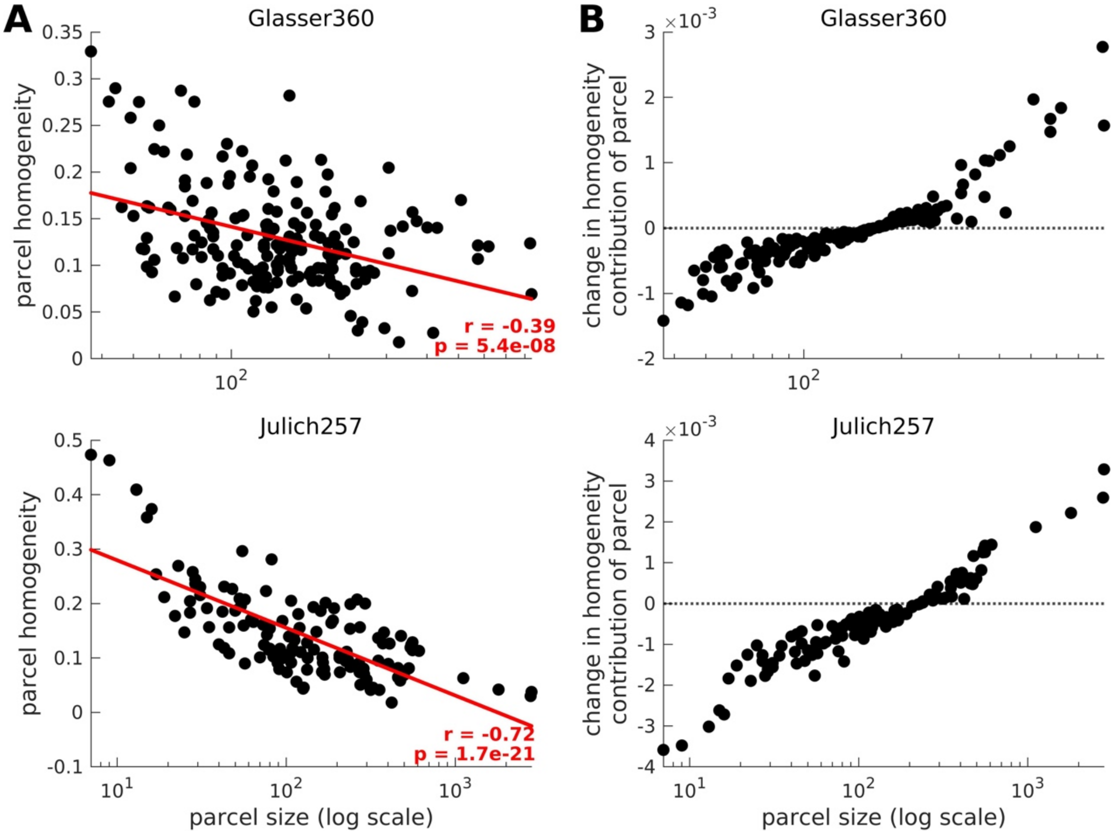

**Supplementary Fig. 13. Relationship between parcel homogeneity and parcel size.** (**A**) Parcel homogeneity as a function of parcel size of the Glasser360 and Julich257 atlases. Parcel homogeneity was based on the FC homogeneity calculated using the HCP resting-state fMRI dataset, and only the left hemisphere results are shown. The dots represent data for each parcel and the red line represents a linear fit with Pearson correlation coefficient $r$ and one-sided p-value $p$. (**B**) Effect of parcel size correction in calculating the overall parcellation homogeneity of the Glasser360 and Julich257 atlases. Here we calculated the change in the contribution of each parcel to the overall homogeneity when weighted averaging was used to correct for parcel size (weights = parcel size) relative to normal averaging (constant weights = 1). The dots represent data for each parcel. Negative/positive values mean that the contribution was decreased/increased. These results demonstrate how our weighted averaging method reduces the homogeneity bias of small parcels.

### S6. Supplementary Tables

**Supplementary Table 1. Existing benchmark parcellations of the human, macaque, marmoset, and mouse neocortices.** The first column shows the species. The second column shows the naming conventions we used in this work, where the number beside the parcellation's name denotes the total number of parcels across the left and right hemispheres. Definitions of the following acronyms are: AAL=Automated anatomical labelling; FVE=Felleman and van Essen; MBM=Marmoset Brain Mapping Project; and ABA=Allen Mouse Brain Atlas. The third column shows the number of parcels in the left (L) and right (R) hemispheres. The last column shows the modalities or features the parcellations were derived from and relevant references.

| Species | Parcellation | Number of parcels (L, R) | Modalities or features used |
|---|---|---|---|
| Human | Brodmann78 | 39, 39 | Cytoarchitecture[30] |
| | Smith88 | 44, 44 | Cyto- and myeloarchitecture[31] |
| | Flechsig92 | 46, 46 | Myelogenesis[32] |
| | Kleist98 | 49, 49 | Lesion-based functionality[33] |
| | Julich257 | 128, 129 | Cytoarchitecture[27] |
| | Desikan70 | 35, 35 | Gyral landmarks[34] |
| | AAL82 | 41, 41 | Sulcal landmarks[35] |
| | Mars82 | 41, 41 | Sulcal landmarks[36] |
| | HarvardOxford96 | 48, 48 | Gyral landmarks[37] |
| | Destrieux150 | 75, 75 | Gyral and sulcal landmarks[38] |
| | Brainnetome210 | 105, 105 | Desikan + functional and diffusion MRI[39] |
| | Cammoun219 | 111, 108 | Desikan + diffusion MRI[40] |
| | Glasser360 | 180, 180 | T1w, T2w, resting-state functional, and task-based functional MRI[28] |
| | Shen200 | 102, 98 | Resting-state fMRI[41] |
| | Craddock271 | 130, 141 | Resting-state fMRI[42] |
| | Schaefer300 | 150, 150 | Resting-state fMRI[43] |
| | Aicha332 | 166, 166 | Resting-state fMRI[44] |
| Macaque | BoninBailey52 | 26, 26 | Cytoarchitecture[45] |
| | Brodmann58 | 29, 29 | Cytoarchitecture[46] |
| | Brodmann82 | 41, 41 | Human cytoarchitectonic parcellation[30] mapped onto the macaque cortex[47] |
| | FVE153 | 77, 76 | Myeloarchitecture and tract-tracing[48] |
| | Markov158 | 80, 78 | Cytoarchitecture[49] |
| | Markov182 | 91, 91 | Cytoarchitecture and immunocytochemistry[50] |
| | Composite233 | 116, 117 | Cyto- and myeloarchitecture, retinotopy, and immunocytochemistry[51] |
| | Saleem266 | 131, 135 | Histology and mean apparent propagator MRI[52] |
| | Paxinos285 | 142, 143 | Cytoarchitecture and immunocytochemistry[53] |
| | Glasser360 | 180, 180 | Human multimodal MRI parcellation[28] mapped onto the macaque cortex[47] |
| Marmoset | MBM108 | 54, 54 | T2w and diffusion MRI and magnetization transfer imaging[54] |

| | | | |
|---|---|---|---|
| | MBM212 | 106, 106 | T2w and diffusion MRI and magnetization transfer imaging[54] |
| | Paxinos228 | 114, 114 | Cytoarchitecture and tract-tracing[54,55] |
| | Riken229 | 114, 115 | Nissl histology and T2w MRI[56] |
| Mouse | ABA80 | 40, 40 | Nissl histology and tract-tracing[26,57,58] |

**Supplementary Table 2. Existing benchmark parcellations of 7 human non-neocortical structures.** The first column shows the structure, where HIP=hippocampus, AMY=amygdala, THA=thalamus, NAc=nucleus accumbens, GP=globus pallidus, PUT=putamen, and CAU=caudate. The second column shows the naming conventions we used in this work, where the number beside the parcellation's name denotes the total number of parcels across the left and right hemispheres. PD=Parkinson's Disease cohort. The third column shows the number of parcels in the left (L) and right (R) hemispheres. The last column shows the modalities or features the parcellations were derived from and relevant references.

| Structure | Parcellation | Number of parcels (L, R) | Modalities or features used |
|---|---|---|---|
| HIP | Melbourne4 | 2, 2 | Resting-state fMRI[59] |
| | Melbourne8 | 4, 4 | |
| | Melbourne10 | 5, 5 | |
| | Julich6 | 3, 3 | Cytoarchitecture[27] |
| AMY | Melbourne4 | 2, 2 | Resting-state fMRI[59] |
| | Julich6 | 3, 3 | Cytoarchitecture[27] |
| THA | Melbourne4 | 2, 2 | Resting-state fMRI[59] |
| | Melbourne8 | 4, 4 | |
| | Melbourne14 | 7, 7 | |
| | Melbourne16 | 8, 8 | |
| | Morel61 | 31, 30 | Cyto- and myeloarchitecture and immunocytochemistry[60] |
| | Oxford13 | 7, 6 | Diffusion MRI[61] |
| NAc | Melbourne4 | 2, 2 | Resting-state fMRI[59] |
| | Cartmell4 | 2, 2 | Diffusion MRI[62] |
| GP | Melbourne4 | 2, 2 | Resting-state fMRI[59] |
| | PD4 | 2, 2 | T1w and T2w MRI[63] |
| PUT | Melbourne4 | 2, 2 | Resting-state fMRI[59] |
| | Melbourne8 | 4, 4 | |
| CAU | Melbourne4 | 2, 2 | Resting-state fMRI[59] |
| | Melbourne8 | 4, 4 | |

**Supplementary Table 3. HCP task contrasts across 7 task types.** The first column shows the task type. The second column shows the number of contrasts within each task type. The last column shows the specific contrast names within each task type.

| Task type | Number of contrasts | Contrasts |
|---|---|---|
| social | 3 | random; tom; tom_random |
| motor | 13 | cue; lf; lh; rf; rh; t; avg; lf_avg; lh_avg; rf_avg; rh_avg; t_avg; cue_avg |

| | | |
|---|---|---|
| gambling | 3 | punish; reward; punish_reward |
| working memory | 19 | 2bk_body; 2bk_face; 2bk_place; 2bk_tool; 0bk_body; 0bk_face; 0bk_place; 0bk_tool; 2bk; 0bk; body; face; place; tool; body_avg; face_avg; place_avg; tool_avg; 2bk_0bk |
| language | 3 | math; story; math_story |
| emotion | 3 | faces; shapes; faces_shapes |
| relational | 3 | match; rel; match_rel |

**Supplementary Table 4. Neurosynth terms divided into 11 cognitive and behavior concepts.** The first column shows the concept. The second column shows the number of terms included within each concept. The last column shows the specific Neurosynth terms included within each concept.

| Concept | Number of terms | Terms |
|---|---|---|
| action | 4 | action; motor control; movement; response selection |
| attention | 8 | attention; distraction; fixation; focus; selective attention; spatial attention; sustained attention; visual attention |
| emotion | 11 | Anxiety; arousal; emotions; emotion regulation; empathy; facial expression; fear; mood; pain; stress; valence |
| executive and cognitive control | 9 | cognitive control; goal; inhibition; maintenance; manipulation; monitoring; planning; response inhibition; updating |
| language | 11 | language; language comprehension; morphology; naming; reading; semantic memory; sentence; speech perception; speech production; verbal fluency; word recognition |
| learning and memory | 20 | adaptation; autobiographical; consolidation; encoding; episodic memory; expertise; face recognition; familiarity; learning; memory; memory retrieval; priming; recall; recognition; rehearsal; reinforcement; retention; retrieval; rule; working memory |
| motivation | 2 | expectancy; task difficulty |
| perception | 11 | detection; discrimination; imagery; integration; mental imagery; multisensory; navigation; object recognition; perception; rhythm; visual perception |
| reasoning and decision making | 12 | categorization; decision; decision making; induction; inference; intelligence; judgement; knowledge; reasoning; reward anticipation; risk; uncertainty |
| social function | 2 | communication; social cognition |
| others | 26 | addiction; anticipation; association; balance; belief; competition; concept; consciousness; context; coordination; eating; efficiency; effort; extinction; gaze; hyperactivity; impulsivity; intention; interference; listening; loss; psychosis; salience; sleep; strategy; thought |

**Supplementary Table 5. Brain-related mouse genes divided into receptor sub-units and interneuron cell-type markers, with grouping types following refs.[57,64].** The first column shows the type of gene. The second column shows the common name of the gene group within each type. The third column shows the number of genes included within each group. The last column shows the specific name of the gene within each group.

| Type | Name | Number of genes | Genes |
|---|---|---|---|
| Receptor | Adrenergic receptor | 5 | *Adra1a*; *Adra1d*; *Adra2a*; *Adra2b*; *Adrb1* |
| | Adenosine receptor | 1 | *Adora2a* |
| | Cannabinoid receptor | 2 | *Cnr1*; *Cnr2* |
| | Cholinergic receptor | 5 | *Chrm1*; *Chrm2*; *Chrm3*; *Chrm4*; *Chrm5* |
| | Dopamine receptor | 4 | *Drd1*; *Drd2*; *Drd3*; *Drd4* |
| | GABA receptor | 1 | *Gabbr2* |
| | Galanin receptor | 2 | *Galr1*; *Galr2* |
| | Glutamate receptor - AMPA | 4 | *Gria1*; *Gria2*; *Gria3*; *Gria4* |
| | Glutamate receptor - Kainate | 5 | *Grik1*; *Grik2*; *Grik3*; *Grik4*; *Grik5* |
| | Glutamate receptor - NMDA | 7 | *Grin1*; *Grin2a*; *Grin2b*; *Grin2c*; *Grin2d*; *Grin3a*; *Grin3b* |
| | Glutamate receptor - Metabotropic | 6 | *Grim1*; *Grim2*; *Grim3*; *Grim4*; *Grim5*; *Grim8* |
| | Histamine receptor | 3 | *Hrh1*; *Hrh2*; *Hrh3* |
| | Hypocretin receptor | 2 | *Hcrtr1*; *Hcrtr2* |
| | Melanocortin receptor | 2 | *Mc3r*; Mc4r |
| | Melatonin-concentrating hormone receptor | 1 | *Mchr1* |
| | Neuropeptide Y receptor | 1 | *Npy1r* |
| | Neurotensin receptor | 1 | *Nrsr1* |
| | Nociceptin receptor | 1 | *Oprl1* |
| | Opioid receptor | 3 | *Oprd1*; *Oprk1*; *Oprm1* |
| | Oxytosin receptor | 1 | *Oxtr* |
| | Purine receptor | 5 | *P2rx1*; *P2ry12*; *P2ry14*; *P2ry2*; *P2ry6* |
| | Serotonin receptor | 8 | *Htr1a*; *Htr1b*; *Htr2b*; *Htr2c*; *Htr3a*; *Htr3b*; *Htr4*; *Htr5b* |
| | Somatostatin receptor | 2 | *Sstr2*; *Sstr4* |
| | Tachykinin receptor | 2 | *Tacr1*; *Tacr3* |
| | Thyrotropin releasing hormone receptor | 1 | *Trhr* |
| | Vasopressin receptor | 2 | *Avpr1a*; *Avpr1b* |
| | VIP receptor | 1 | *Vipr2* |
| Cell-type marker | Parvalbumin | 1 | *Pvalb* |
| | Somatostatin | 1 | *Sst* |
| | Calbindin | 2 | *Calb1*; *Calb2* |
| | Vasoactive intestinal polypeptide | 1 | *Vip* |
| | Myelin marker | 2 | *Mbp*; *Plekhb1* |
